# Cell-free DNA fragmentomic characteristics in transposon elements inform molecular regulators and enhance cancer diagnosis

**DOI:** 10.1101/2025.07.10.664250

**Authors:** Fanglei Gong, Yuqi Pan, Huizhen Lin, Yunyun An, Mengqi Yang, Xiaoyi Liu, Yunxia Bai, Wanqiu Wang, Zhenyu Zhang, Bianbian Tang, Kun Zhang, Xin Zhao, Yu Zhao, Changzheng Du, Xuetong Shen, Kun Sun

## Abstract

Fragmentomics of plasma cell-free DNA (cfDNA) are emerging diagnostic biomarkers in cancer liquid biopsy, while the molecular regulations of cfDNA fragmentation remain elusive. In this study, we investigated the cfDNA fragmentomics in transposon elements (TEs), a special category of sequences accounting for around half of human genome. We discovered dynamics in fragmentomic features across various types of TEs in human cfDNA, including size, ending patterns and coverage, which were validated in mice and dogs. These dynamics highly correlate with epigenomic features associated with chromatin states, including DNA methylation and histone modification signals, demonstrating fundamental regulatory roles of chromatin state in cfDNA production. Furthermore, fragmentomic features within TEs were significantly altered in cancer samples, offering improved diagnostic performance compared to genome-wide fragmentomic measurements. Additionally, cfDNA coverage in TEs showed frequent imbalances between cancer and control samples in a TE- and cancer type-dependent manner, presenting a promising biomarker for cancer diagnosis. Leveraging artificial intelligence (AI) on cfDNA fragmentomic features within TEs, we developed high-performance models, named TEANA (<u>TE A</u>nalysis in cell-free D<u>NA</u>), for cancer diagnosis and tumor-origin prediction. The models were validated across two pan-cancer datasets with more than 1600 samples. Hence, dynamics of cfDNA fragmentomics within TEs, which is highly correlated with cfDNA epigenomics, shed light on the regulatory mechanism of cfDNA biology and highlight translational potential in cancer liquid biopsy.

## Introduction

Circulating cell-free DNA (cfDNA) in plasma has been a subject of research for over 75 years ^1^. In healthy subjects, cfDNA predominantly originate from the hematopoietic system and liver ^2,3^, whereas in cancer patients, tumors release cfDNA into the plasma ^4^. The analysis of cfDNA to assess the presence, progression, and molecular characteristics of tumors is commonly referred to as “cancer liquid biopsy”, a field that has seen significant advancements in translational medicine over the past decade ^5^. Topologically, most cfDNA molecules are small fragments, typically under 200 base pairs (bp), generated during cell death ^6^. Recent studies have proven that the generation of fragmented cfDNA molecules is not a random process ^7–11^. Distinctive features include a peak at 166 bp and a 10 bp periodicity below 143 bp ^12,13^, as well as biased end sequence usages ^14^. The fragmentation patterns of cfDNA, referred to as “fragmentomics”, are frequently altered in cancer, making them promising biomarkers for cancer diagnosis and tumor origin tracing ^15–19^. However, despite the substantial technological progresses in cfDNA analysis, molecular mechanisms regulating cfDNA fragmentation is still much less elucidated. In a previous study, we have demonstrated that DNA methylation plays a crucial role by influencing the cleavage preferences of the DNASE1L3 endonuclease (which is associated with CCCA end motif usage in cfDNA) ^13,14,20^. Recent studies reported that H3K27ac histone modification and gene expression levels are also associated with cfDNA fragmentation ^8,21,22^. However, given that these epigenomic modifications and genes only constitute a small proportion of the human genome, it suggests the existence of additional molecular regulators or mechanisms for cfDNA fragmentation that remain to be explored.

In cancer biology, transposon elements (TEs) are known to undergo frequent dysregulations. TEs, often referred to as “mobile DNA” or “jumping genes”, play a significant role in shaping the genetic landscape and account for approximately half of the human genome ^23^. TE activity is regulated by epigenomic factors (e.g., DNA methylation and histone modifications) and is predominantly silenced in normal cells; however, in malignant cells, epigenomic remodeling and dysregulation can activate TEs in a cancer-and TE-type-dependent manner ^24,25^. The presence or activity of TEs in tissues has been widely utilized as biomarkers for cancer diagnosis and prognosis ^26,27^. Moreover, alterations in TE-associated tumor-derived cfDNA have been investigated, particularly concerning their copy number or DNA methylation patterns ^28,29^. For instance, Grabuschnig et al. reported that TEs are overrepresented in plasma cfDNA, with coverage patterns shift in sepsis ^30^. Douville et al. developed a PCR-based assay called RealSeqS (Repetitive Element AneupLoidy Sequencing System), which detected aneuploidy in ∼50% blood samples ^31,32^. Additionally, Annapragada et al. introduced the ARTEMIS (Analysis of RepeaT EleMents in dISease) and ARTEMIS-DELFI models that utilized machine learning on sequence features of TEs, demonstrating promising performance in cancer diagnosis and tumor origin prediction ^33,34^. Collectively, these studies highlight the potential of TEs in cancer liquid biopsy; however, the characteristics of cfDNA in TEs, particularly fragmentomic features and dynamics in cancer patients, remain largely unexplored and warrant further investigations. Therefore, in this study, we investigate cfDNA fragmentomic features in TEs and their relationships with various epigenomic factors to uncover the underlying molecular regulators of cfDNA fragmentation. Furthermore, translational potential of cfDNA fragmentomic dynamics in TEs for cancer liquid biopsy is also explored.

## Results

### Fragmentomic characteristics of transposon elements in cfDNA

Considering that most studies generate low-pass cfDNA data, we ranked the TEs by their copy numbers and picked up those with more than 10,000 copies to avoid stochastic variations in the low-copy TEs. As a result, 16 family of TEs showed up, which includes 11 retrotransposons covering the common SINE (ALU and MIR), LINE (L1, L2, CR1, and RTE-X), and LTR (ERVL-MaLR, ERV1, ERVL, Gypsy, and ERVK) classes, as well as 5 DNA transposons (hAT-Charlie, TcMar-Tigger, hAT-Tip100, hAT-Blackjack, and TcMac-Mariner; Suppl. Table S1). Together, these 16 TEs account for ∼45% of human genome and were kept for downstream analyses.

We first explored the widely studied cfDNA fragmentomic characteristics (size and end motif), in these transposon elements using a cohort of 24 healthy subjects ^13^. We found that cfDNA in 15 out of 16 TEs were significantly shorter than genome-wide level, as evidenced by increased proportions of short fragments (i.e., ≤150 bp); in the meantime, the size distributions also varied across TEs (Fig. 1a). The size distributions of representative TEs in one sample were illustrated in Fig. 1b. For end motif patterns, compared to genome-wide level, the CCCA end motif usages were significantly elevated in 4 while decreased in 10 TEs, and not changed in the rest 2 TEs; moreover, the diversities of end motif usages were significantly elevated in 2 while decreased in the rest 14 TEs (Fig. 1a). Of note, among all TEs, the SINE/ALU retrotransposon showed the highest CCCA end motif usage and lowest motif diversity. In addition, all TEs showed significantly altered sequencing depth in relative to genome-wide level (defined as RSD, and is always 1 for genome-wide level; see Methods; Fig. 1a) with high variations across different types of TEs. For instance, LTR/ERVL-MaLR was 35% higher than DNA/TcMar-Mariner.

**Fig. 1.**
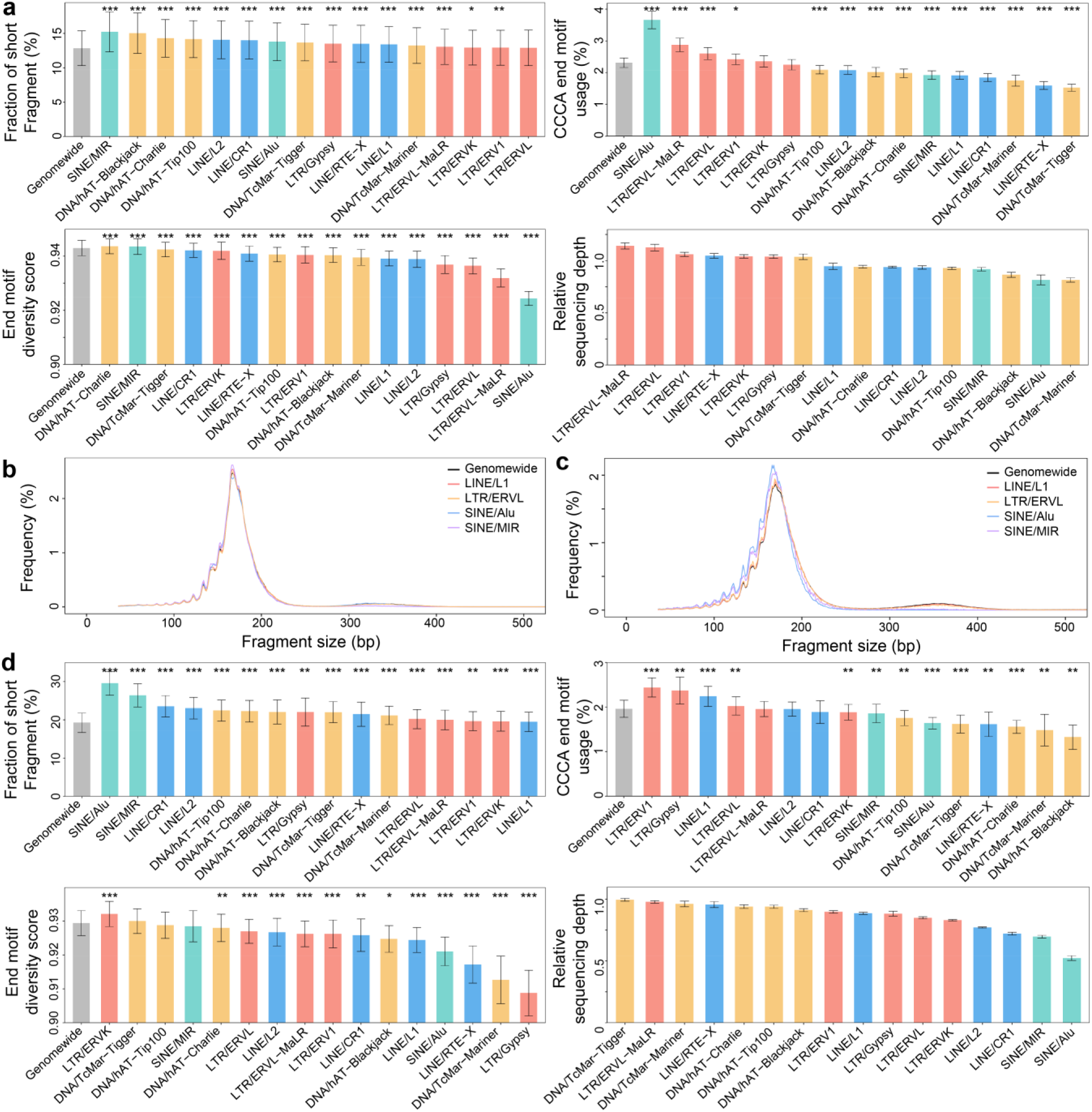
CfDNA fragmentomic characteristics in transposon elements (TEs). (a) CfDNA size, motif patterns, and relative sequencing depths across 16 TEs in 24 human samples. (b-c) Size distributions of cfDNA in representative transposon elements in (b) human and (c) murine samples. (d) CfDNA size, motif patterns, and relative sequencing depths across 16 TEs in 6 murine samples. In (a,d), bars indicated mean ± s.d., and p-values were calculated between each TE and genome-wide level using paired t-tests and left blank for p>0.05. *p<0.05; **p<0.01, ***p<0.001.

To validate the findings, we profiled the cfDNA fragmentomics in 16 and 14 most abundant TEs in mouse (N=6; generated in this study) and dog (N=9; collected from Favaro et al. study ^35^) plasma samples, respectively (Suppl. Table S1). Of note, the high-abundant TEs were different across these 3 species, which might be associated with their appearances during mammalian evolution ^36^. The results on cfDNA fragmentomics were shown in Fig. 1c-d and Suppl. Fig. S1. Overall, the patterns were highly consistent with human data that cfDNA fragmentomics were frequently altered in TEs, while the characteristics could be different among species. For example, SINE/ALU showed significantly decreased CCCA end motif patterns in mice, while this TE was too rare to be included in the current analysis in dogs; LTR/Gypsy showed higher CCCA end motif usages than genome-wide level in both mice in dogs, while not in human. Together, these results revealed variable fragmentomic characteristics of cfDNA among TEs. Given that the DNA sequences are highly similar with all the copies of the same type of TE, these elements might serve as a unique material to explore the regulators and biology of cfDNA fragmentation.

### Regulatory roles of epigenomics on cfDNA fragmentomics

To explore the molecular regulators affecting cfDNA fragmentomics in TEs, we correlated the fragmentomic features with various epigenomic regulators. Firstly, we investigated DNA methylation, which is a known genome-wide regulator of cfDNA fragmentation. Hyper-methylation is a known mechanism in silencing TEs ^37^, while we still observe a remarkable number of TE copies with hypo- or partial-methylation (Suppl. Table S2). The results for SINE/ALU and LTR/ERVK were illustrated as examples in Fig. 2, and the others could be found in Suppl. Fig. S2-S8. In all TEs, hypo-methylated copies showed significantly elevated fractions of short cfDNA fragments compared to partial- and hyper-methylated ones, as well as significantly decreased sequencing depths; in most TEs, hypo-methylation was also associated with altered CCCA end motif usages and overall end motif diversities, while the directions vary across TEs. Interestingly, in many TEs, partially methylated copies did not always show intermediate fragmentomic characteristics between hypo- and hyper-methylated ones; for instance, in both SINE/ALU and LTR/ERVK, partially methylated copies showed even lower fraction of short fragments than both hypo- and hyper-methylated ones, suggesting the presence of additional regulators besides DNA methylation.

**Fig. 2.**
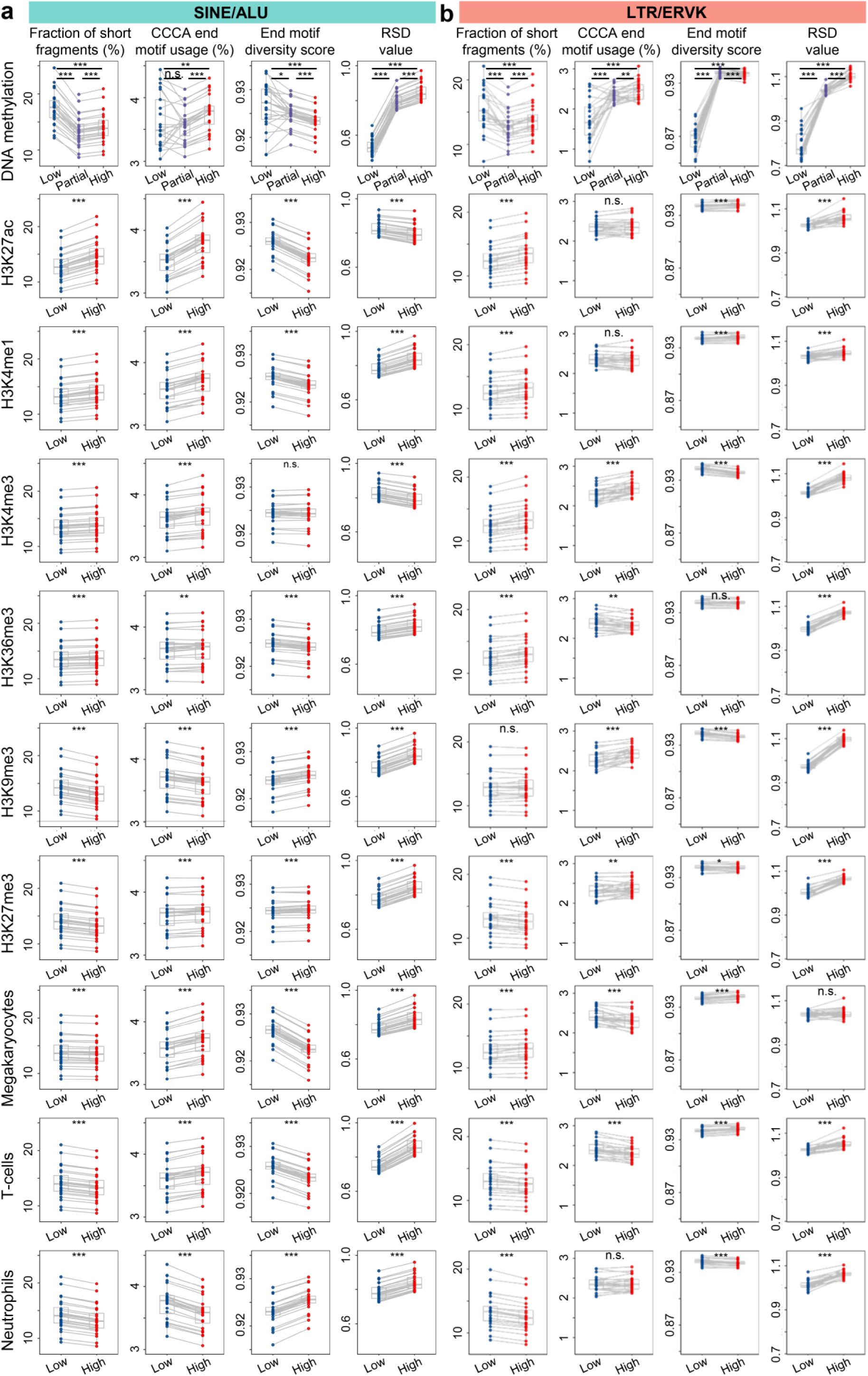
CfDNA fragmentomic characteristics in (a) SINE/ALU and (b) LTR/ERVK in different epigenomic context. For DNA methylation analysis, copies in SINE/ALU and LTR/ERVK were divided into three groups based on the average DNA methylation levels of CpG sites they covered; for histone modification (H3K27ac, H3K4me1, H3K4me3, H3K36me3, H3K9me3, and H3K27me3) and open chromatin (megakaryocyte, T-cells, and neutrophils) analysis, copies in SINE/ALU and LTR/ERVK were divided into two groups based on the size-normalized signals of these epigenomic markers. P-values were computed using paired t-tests. *p<0.05, **p<0.01, ***p<0.001.

To explore the potential regulators, we investigated 6 fundamental histone modifications (i.e., H3K27ac, H3K4me1, H3K4me3, H3K36me3, H3K9me3, and H3K27me3) in cfDNA assayed with cfChIP-seq experiments ^38,39^. For each histone modification, we split each family of TE copies into 2 equal-sized groups (i.e., “low” and “high”) based on the signals of this histone modification, then we compare the fragmentomic characteristics between these two groups. Overall, we observed significant differences in various fragmentomic characteristics between “low” and “high” groups in a TE- and histone modification-dependent manner (Fig. 2 and Suppl. Figure S2-S8). Of note, for almost all TEs, copies with lower signals in active chromatin-related histone modifications (H3K27ac and H3K4me1 for enhancers, H3K4me3 for promoters, and H3K36me3 for transcribed gene bodies) showed lower fractions of short fragments, while those with lower signals in histone modifications associated with repressive or heterochromatin (H3K9me3 and H3K27me3) showed opposite trends. We further investigated the effect of chromatin accessibility in 3 types of hematopoietic cells that are the known major contributors of cfDNA ^2,3,38,40^: megakaryocytes, T-cells, and neutrophils. Again, we observed significant differences in various fragmentomic characteristics between TE copies with “low” and “high” chromatin accessibility signals in these cells. Interestingly, in most TEs, the differences in fragmentomic characteristics were consistent between megakaryocytes and T-cells while not in neutrophils, which might be associated with different cell death scheme in neutrophils, e.g., through neutrophil extracellular traps ^41,42^, and might be worthwhile for further investigations. Together, the results suggested that besides DNA methylation, histone modifications and chromatin accessibility might also possess regulatory roles on cfDNA fragmentomics.

As transposable elements (TEs) constitute approximately half of the human genome, we sought to extend the analysis to explore how epigenomic modifications influence cfDNA fragmentation across the entire genome. To this end, we utilized the signals of 6 histone modifications in cfDNA and segmented the genome into 15 chromatin states (Fig. 3a and Suppl. Table S3) ^43,44^. This analysis revealed significant differences in cfDNA fragmentomics across chromatin states in the 24 non-cancerous human samples (Fig. 3b-c). Specifically, cfDNA molecules in active chromatin states (such as promoters, enhancers, and transcribed regions) were generally shorter with lower depth, while those in repressive chromatin states (including heterochromatin and polycomb-repressed regions) exhibited the opposite pattern. Additionally, regions with strong transcriptional activity (indicated by higher H3K36me3 signals) showed distinct fragmentation patterns compared to regions with weaker transcription (indicated by lower H3K36me3 signals), indicating the relationship between cfDNA fragmentomics and gene expression ^21^; the motif patterns were highly variable across chromatin states, suggesting dynamic participation of endonucleases in cfDNA fragmentation ^45^.

**Fig. 3.**
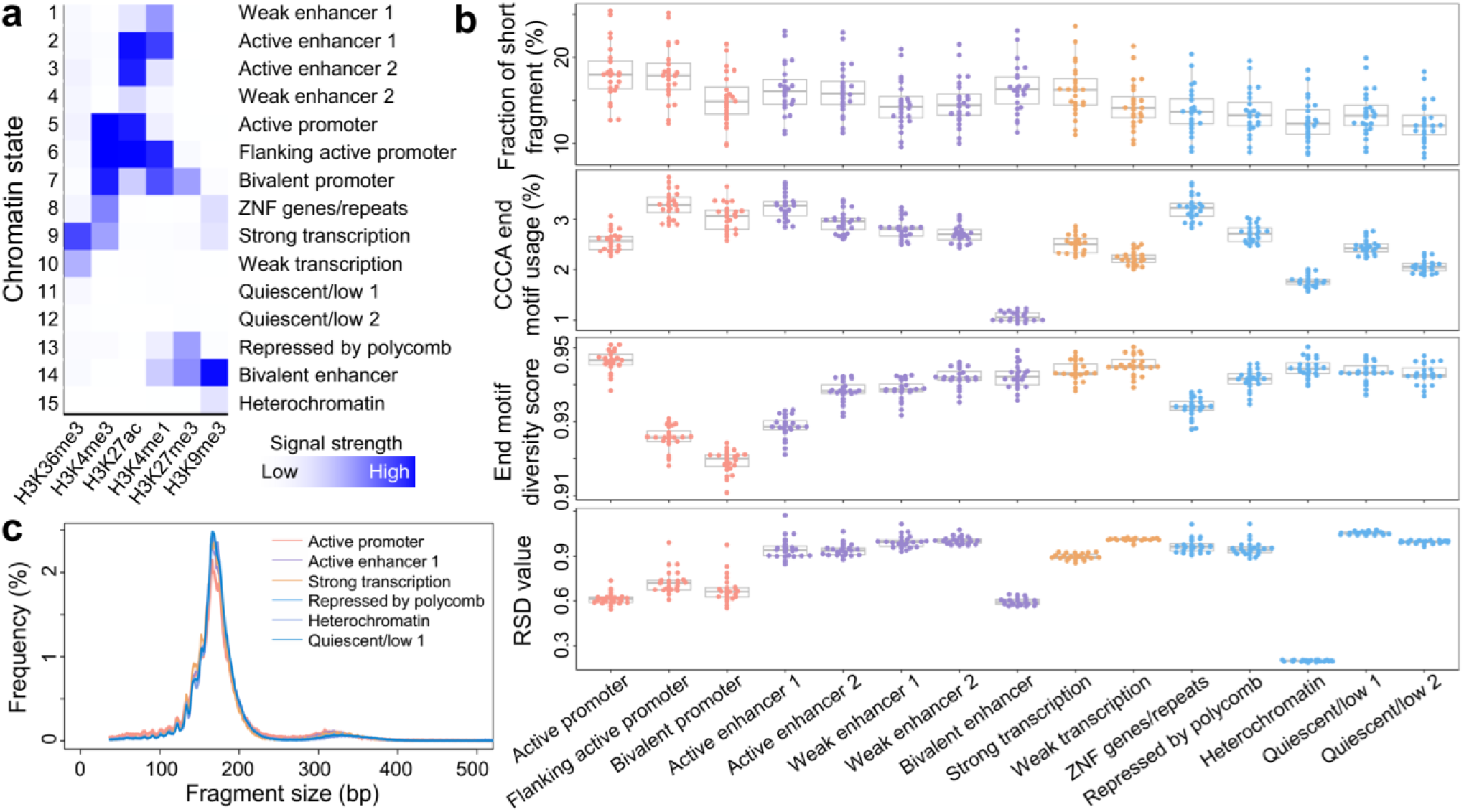
CfDNA fragmentomic characteristics across different chromatin states in human genome. (a) Mapping and annotation of chromatin states (N=15) using 6 histone modification signals in cfDNA; cfDNA size, motif patterns, and relative sequencing depth (RSD) across different chromatin states in a cohort of 24 control subjects; and (c) cfDNA size distribution of representative chromatin states in one control subject. In b, for all features, P<10^-10^ across chromatin states calculated by one-way repeated-measures ANOVA.

### CfDNA fragmentomic characteristics in TEs in Hepatocellular Carcinoma

Considering that TEs are frequently activated by epigenomic dysregulations in cancer, we wonder whether the fragmentomic features of TEs in cfDNA would alter in cancer patients. To this end, we further investigated the fragmentomic patterns of cfDNA in TEs in a cohort of 56 pre-treatment Hepatocellular Carcinoma patients (HCC; 31, 10, and 15 were in stage A, B, and C, respectively, according to the Barcelona Clinic Liver Cancer staging system) and 24 age- and gender-matched non-cancerous controls ^46^. The average sequencing depth in this cohort was ∼3x haploid human genome coverage. As expected, cfDNA size and end motif patterns in TEs were altered in HCC patients in the same direction as the genome-wide level and showed comparable performance in differentiating HCC patients from controls (Fig. 4a and Suppl. Fig. S9). However, in 12 TEs, the RSD values showed significant differences between HCC patients and controls (Fig. 4b); Receiver Operating Characteristic (ROC) analysis showed that RSD values in these 12 TEs could readily differentiate HCC patients from controls with Area Under the ROC Curves (AUCs) ranged from 0.65 to 0.91 (all P<0.05, Z-tests; Fig. 4c). Another cfDNA fragmentomic feature, E-index, which measures the consistency in the cfDNA ends between a sample-of-interest to a pool of healthy controls ^13^, showed a decrease trend in HCC patients in genome-wide level while did not reach statistical significance (P=0.27, Mann-Whitney U test; Fig. 4d); interestingly, for both HCC patients and controls, E-index values declined in all TEs compared to genome-wide level (all P<10^-5^, paired t-tests), while the reductions were more profound in HCC patients such that the E-index values in all TEs become significantly lower than that in controls (Fig. 4d), which allowed us to perform cancer diagnosis (Fig. 4e). Together, these results demonstrated that the cfDNA fragmentomic patterns in TEs were altered in HCC patients and could be informative in cancer diagnosis.

**Fig. 4.**
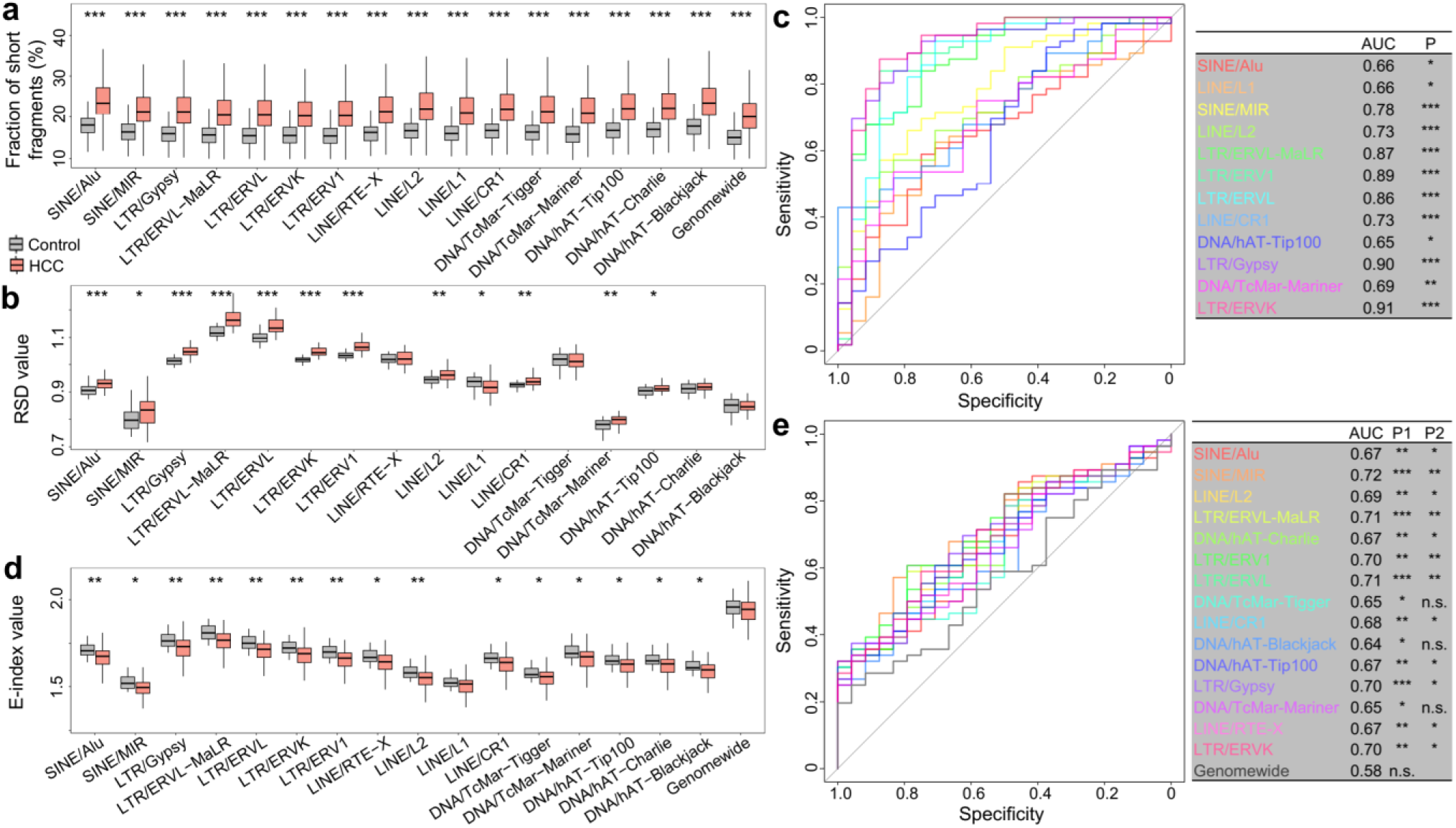
CfDNA fragmentomic characteristics in transposon elements (TEs) in hepatocellular carcinoma (HCC) patients. (a) Fraction of short cfDNA fragments (i.e., < 150bp) between controls and HCC patients across various TEs. (b) Relative sequencing depth (RSD) values between controls and HCC patients across various TEs and (c) corresponding Receiver Operating Characteristic (ROC) curves for diagnosis. (d) E-index values between controls and HCC patients across various TEs and (e) corresponding ROC curves for diagnosis. In (a,b,d), grey and orange boxes presented controls and HCC samples, respectively; p-values were calculated using Mann-Whitney U tests and left blank if p>0.05. In (c,e), the TEs were limited to those showing significant differences between HCC samples and controls in (b, d), respectively; P-values indicated the significances for Area Under the ROC Curves (AUCs) against random assignments, calculated by Z-tests. In (e), P1 was the p-value for AUC calculated using Z-tests, and P2 was the p-value comparing the ROC of each TE versus genome-wide level using DeLong test. *p<0.05, **p<0.01, ***p<0.001.

### CfDNA fragmentomic characteristics in TEs in pan-cancer

With the inspiring findings in HCC, we then explored cfDNA fragmentomic features in TEs in other cancer types. Firstly, we investigated a dataset from Liang et al. ^47^ which contained 10 HCC patients, 8 lung cancer samples (after removing 2 post-treatment samples), and 10 controls. As shown in Fig. 5a-b and Suppl. Fig. S10, highly consistent results to our HCC cohort were observed. In particular, in various TEs, the RSD values showed significant differences between cancer patients and controls, which enabled cancer diagnosis with AUCs ranging from 0.79 to 0.98 (all P<0.05, Z-tests; Fig. 5b and Suppl. Fig. S10b). Interestingly, the patterns were not identical between HCC and lung cancer. For example, among the 5 DNA transposons, TcMar-Tigger and TcMar-Mariner showed more profound elevations in lung cancer samples, while the elevations were more remarkable in hAT-Tip100, hAT-Charlie, and hAT-Blackjack in HCC samples (Fig. 5a). This result suggested cancer type specificities of cfDNA fragmentomic characteristics across TEs, which might be informative in tumor-origin tracing.

**Fig. 5.**
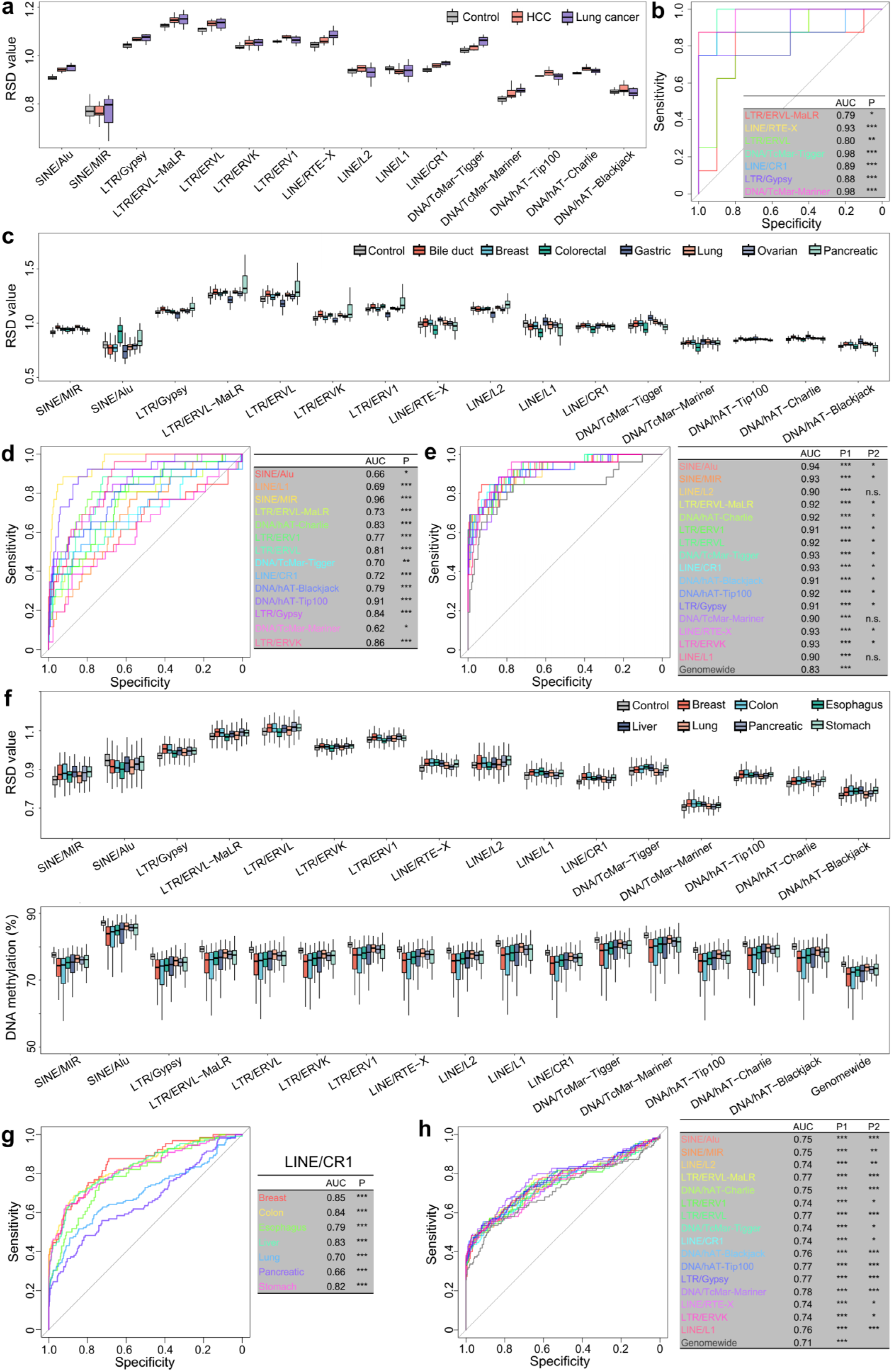
CfDNA fragmentomic characteristic in 16 transposon elements (TEs) in pan-cancer. In Liang et al. dataset, (a) Relative sequencing depth (RSD) values across TEs. Seven TEs showed significantly different RSD values between lung cancer samples and controls (P<0.05, Mann-Whitney U tests), and their ROC curves for cancer diagnosis in lung cancer were shown in (b). In Cristiano et al. dataset, (c) RSD values across TEs. Fourteen TEs showed significantly different RSD values between bile duct cancer samples and controls, and (d) showed the ROC curves for cancer diagnoses in these 14 TEs. (e) ROC curves for cancer diagnoses using E-index values in 16 TEs versus genome-wide level in bile duct cancer. In Bie et al. dataset, (f) RSD values and DNA methylation densities across TEs. (g) ROC curves in cancer diagnoses using RSD values in LINE/CR1 among cancer types, and (h) ROC curves for cancer diagnoses using DNA methylation densities in TEs versus genome-wide level in stomach cancer. In b,d,e,g,h, P and P1 were p-values for AUCs calculated using Z-tests, and P2 was the p-value comparing the ROC of each TE versus genome-wide level using DeLong test. *p<0.05, **p<0.01, ***p<0.001.

Next, we analyzed a pan-cancer dataset from Cristiano et al. ^34^ containing 208 cancer samples from 7 cancer types and 215 controls (Fig. 5c-e and Suppl. Fig. S11-S19). Consistent with our HCC cohort and Liang et al. dataset, the RSD values showed significant differences in cancer samples in various TEs. For instance, RSD values were altered in 14 TEs in bile duct cancer samples (except for LINE/L2 and LINE/RTE-X), and their corresponding AUCs for cancer diagnoses ranged from 0.62 to 0.96 (all P<0.05, Z-tests; Fig. 5d). In addition, in cancer samples, the decreases of E-index values were also larger in most TEs compared to genome-wide level, which resulted in improved performances in cancer diagnoses (Suppl. Fig. S11). Fig. 5e showed the ROC curves for bile duct cancer samples: the AUCs for E-index values in TEs ranged from 0.90 to 0.94, most of which were significantly higher than the genome-wide level’s 0.83 (all P<0.05 except for DNA/TcMar-Mariner, LINE/L1, and LINE/L2, DeLong tests). The results for other cancers could be found in Suppl. Fig. S12-S18. When comparing the fragmentomic features in various TEs for pan-cancer diagnoses, E-index values in SINE/ALU showed the best performances with AUCs ranging from 0.73 in ovarian cancer to 0.94 in bile duct cancer (all P<0.001, Z-tests; Suppl. Fig. S19a), while the RSD values in SINE/MIR showed the best performances with AUCs ranging from 0.74 in pancreatic cancer to 0.98 in lung cancer (all P<0.001, Z-tests; Suppl. Fig. S19b). Moreover, consistent with the observations in Liang et al. dataset, alterations of RSD values also showed remarkable variabilities among cancer types. For example, in SINE/ALU, RSD values were elevated in colorectal and pancreatic cancers, while they were decreased in the other 5 cancer types. Together, the results suggested that aberrated coverages of TEs in cancer samples could be both cancer-and TE-type dependent.

To further validate the findings, another pan-cancer dataset from Bie et al. ^48^ were analyzed, which contained 749 cancer samples covering 7 cancer types and 451 controls after quality control. Highly consistent results to that in Cristiano et al. dataset were observed (Fig. 5f-h and Suppl. Fig. S20-S27). Of note, in all TEs, RSD values showed significant differences between cancer samples and controls, while LINE/CR1 showed the highest performance in cancer diagnosis, resulting in AUCs ranging from 0.66 in pancreatic cancer to 0.85 in breast carcinoma (all P<0.001, Z-tests; Fig. 5g). Notably, this dataset was generated using NEBNext Enzymatic Methyl-seq (EM-seq) protocol ^48,49^, which allowed us to analyze the DNA methylation profiles of TEs along with fragmentomic features. We found that in most cancers, DNA methylation levels in TEs could provide improved diagnostic performance than genome-wide level. For example, in stomach cancer, DNA methylation levels in all 16 TEs produced higher AUCs than genome-wide level for cancer diagnosis (all P<0.05, DeLong tests; Fig. 5h). Together, the results revealed dynamics of cfDNA fragmentomics and epigenomics in TEs in cancers, as well as their favorable performances in diagnoses.

### AI-enhanced models for cancer diagnosis and tumor origin tracing

Given the alterations of various cfDNA fragmentomic characteristics in cancer, we evaluated the feasibility of integrating such fragmentomic characteristics in TEs as features to build high-performance cancer diagnosis models. We named this approach TEANA-Dx (TE analysis in cell-free DNA for cancer diagnostics) and evaluated it on two pan-cancer datasets: Cristiano et al. ^34^ and Bie et al. ^48^ with 423 and 1201 samples, respectively. The performances were assessed using 10-fold cross-validation repeated 10 times such that all the samples were only predicted by models trained without themselves to eliminate information disclosure ^34,46,48^. The results for TEANA-Dx models utilizing different set of features were summarized in Fig. 6 and Table 1. Briefly, in Cristiano et al. dataset, TEANA-Dx models with RSD or E-index feature alone showed AUCs of 0.92 and 0.88, respectively, while it achieved 0.96 when RSD, E-index, size, and CCCA end motif usage were all utilized (all P<10^-10^, Z-tests; Fig. 6a). Moreover, the AUCs ranged from 0.92 to 0.99 across cancer types (Fig. 6b) and achieved 0.92 and 0.96 for stage I and II cancer samples (Fig. 6c), respectively. The AUCs were higher than previous models, including DELFI ^34^, ARTEMIS-DELFI ^33^, and EXCEL ^46^, while TEANA-Dx used much less features (N=80 in TEANA-Dx vs 500-1280 in others). Moreover, TEANA-Dx reached an overall sensitivity of 85.1% at 95% specificity (Fig. 6a and Suppl. Table S4), which was significantly higher than DELFI on the same dataset (P=0.038, Binomial test). Similarly, in Bie et al. dataset (Fig. 6d-f), TEANA-Dx model with RSD, E-index, size, CCCA end motif usage, and DNA methylation levels achieved an overall AUC of 0.95, at least 0.91 for all cancer types and at least 0.89 for all stages (all P<10^-10^, Z-tests), which were comparable to the THEMIS model ^48^ on the same dataset. Moreover, in the original study, Bie et al. split the dataset into non-overlapping training and testing subsets ^48^. We there re-trained the TEANA-Dx model only using the training subset, then directly applied the trained model on the testing subset. As shown in Fig. 6g-i, this TEANA-Dx model still showed high performance on the testing subset with an overall AUC of 0.93, ranging between 0.86-0.96 across cancer types, and 0.87-0.96 across stages (all P<10^-10^, Z-tests).

**Fig. 6.**
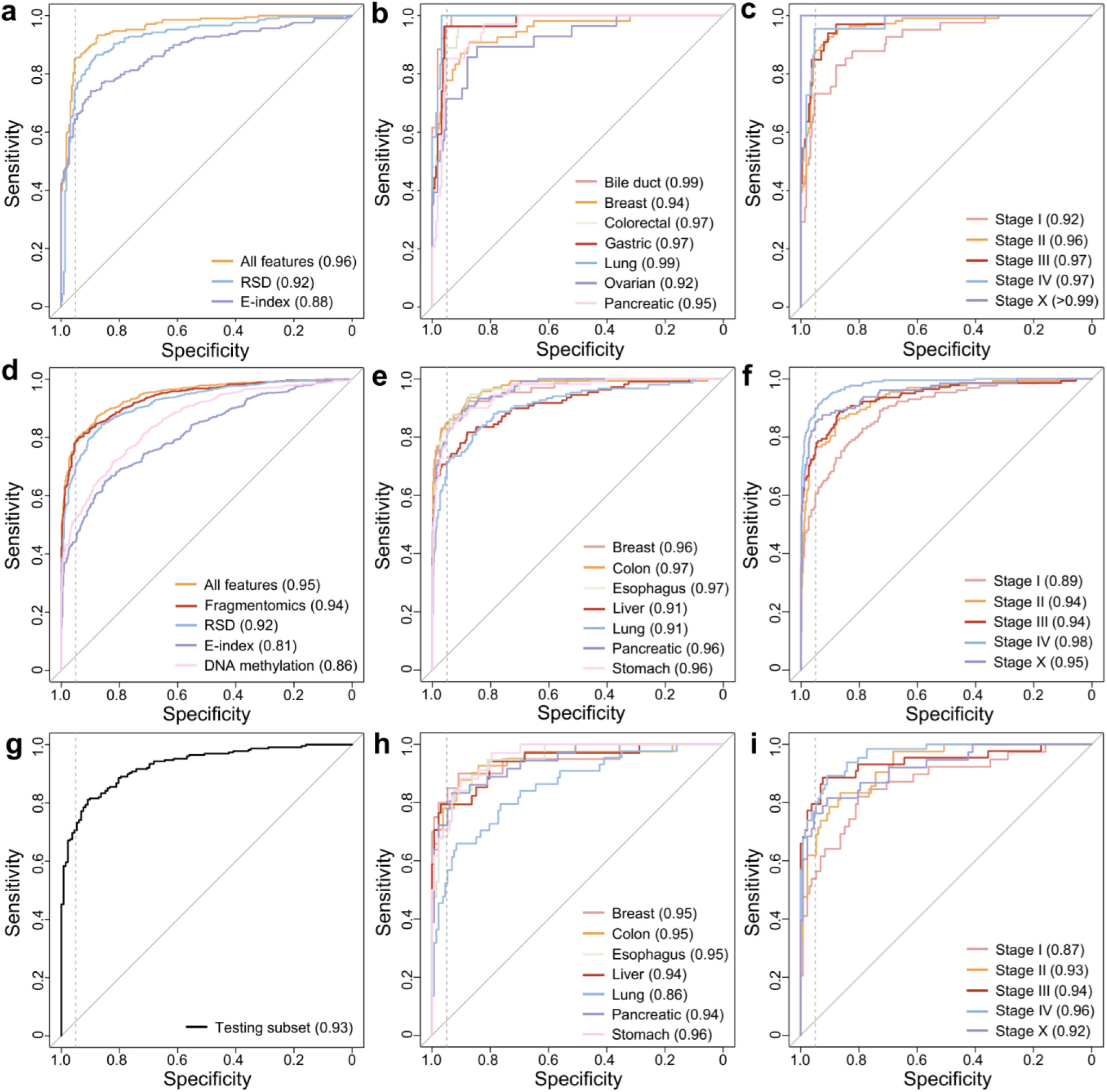
ROC curves of TEANA-Dx models for cancer diagnosis. In Cristiano et al. dataset, ROC curves of (a) TEANA-Dx models using E-index values only, RSD values only, or all features, (b) for each cancer type, and (c) for each stage. In Bie et al. dataset, ROC curves of (d) TEANA-Dx models using E-index values only, RSD values only, DNA methylation only, or all features, (e) for each cancer type, and (f) for each stage. We trained another TEANA-Dx model using the training subset in Bie et al. dataset and directly applied the model on the non-overlapping testing subset, and (g-i) showed the ROC curves for (h) overall samples, (i) across different cancer types, and (i) stages. Dashed lines indicated 95% specificity. The AUCs were recorded in parentheses, and P<10^-10^ for all AUCs, Z-tests.

We further explored the feasibility of using fragmentomic features in TEs to predict the tissue-origin of the tumors in cancer patients, which resulted in a classification model named TEANA-Top (TE analysis in cell-free DNA for tumor-origin prediction). In Cristiano et al. dataset, the top-ranked tissue-origin predictions by TEANA-Top achieved an overall accuracy of 56.7% (118 out of 208), ranged from 25.0% to 67.6% across cancer types, which were significantly higher than random assignment (P<0.05 for all cancer types, Binomial tests; Suppl. Table S5) and comparable to the DELFI and ARTEMIS-DELFI approaches (both P>0.1, Binomial tests) ^34^. Together, the results demonstrated high translational significance of fragmentomic analyses in TEs for cancer liquid biopsy.

## Discussion

In this study, we performed a comprehensive investigation on cfDNA fragmentomics in TEs, the “mobile elements” contributing around half of human genome. In human, mice, and dogs, dynamics in cfDNA fragmentomics across TEs were all observed, which inspired the exploration on associations between various epigenomic regulators and cfDNA fragmentomics. Notably, for each TE family, the genomic sequences of its thousands of copies in the human genome share high similarities, which makes TEs ideal models for studying the regulatory roles of epigenomics on cfDNA fragmentomics as they can minimize the effect of genomic differences in DNA sequences. We first validated that DNA methylation levels and H3K27ac signals affect cfDNA fragmentomics in TEs (Fig. 2), which were consistent with previous studies ^13,22^. More importantly, besides DNA methylation, we further explored the potential roles of various fundamental histone modifications on cfDNA fragmentation patterns. In biology, key histone modifications define chromatin states ^50^, and DNA is packed more tightly in repressive chromatins compared to active ones. Active chromatins are usually associated with hypo-DNA methylation and H3K27ac, H3K4me1, or H3K4me3, while repressive chromatins are associated with hyper-DNA methylation and H3K27me3 or H3K9me3; highly expressed genes are usually marked with elevated DNA methylation levels and H3K36me3 signals in their bodies ^51^. As shown in Fig. 2-3 and Suppl. Fig. S2-S8, active chromatin states are usually associated with shorter size and repressive chromatin states are associated with longer size, while the motif patterns were highly dynamic across the chromatin states, suggesting that chromatin state might implicate the activities of endonucleases and cfDNA size. Together, the findings were consistent with previous studies reporting the relationship between H3K27ac signals, gene expressions and cfDNA fragmentomics^21,22^, and suggested that chromatin states, defined by various histone modifications and DNA methylation, composed a complex network to regulate cfDNA fragmentation. In addition, various previous studies had demonstrated the feasibility and significance of deducing gene expressions from cfDNA using nucleosomal spacing and promoter coverages ^8,9,52–54^; as shown in Fig. 3b, the drastically different cfDNA fragmentation patterns between strong and weak transcription regions suggested that cfDNA fragmentomic characteristics in gene bodies might also be informative for this task therefore deserve further explorations.

Besides explorations on the biology of cfDNA fragmentomics, we also found that some biomarkers (e.g., E-index and DNA methylation) showed improved diagnostic performances on TEs than genome-wide level (Fig. 4-5), which is consistent with previous reports that cfDNA favor fragmentation hotspots in TEs ^55^. In fact, in non-malignant somatic cells, most TEs are repressed through epigenetic mechanisms; while in malignant cells, TEs are frequently activated due to global epigenetic dysregulations (including de-methylation), which may lead to increased aberrations in cfDNA fragmentomic characteristics. Besides size and end motif patterns, the relative sequencing depth (RSD) of various TEs in cfDNA showed significant differences in cancer samples compared to controls, which parameter could serve as a novel biomarker for cancer diagnosis. Of note, as validated in two large-scale pan-cancer datasets, the SINE/MIR (mammalian-wide interspersed repeat) family demonstrates greater aberration and superior diagnostic performance than other TEs. In fact, SINE/MIRs are the most ancient family of TEs in human and the only TE that could serve as enhancers/promoters to regulate tissue-specific gene expression, and they are also extensively activated in cancer ^56–58^.

In addition, with AI, we developed high-performance models, TEANA, for cancer diagnosis and tumor-origin prediction, as validated in two large-scale pan-cancer cohorts. In fact, individual features are not perfect for diagnosis and might not always work well across datasets. For example, RSD outperforms other fragmentomic features (e.g., E-index) in our HCC cohort, while E-index showed better performance than RSD in Cristiano et al. and Bie et al. datasets. The underline reasons are unclear and might be related with the experimental protocols and batch effects, while highlights the advantage of stacked diagnostic models that integrate multiple features towards enhanced and more stable performances. Although TEANA and ARTEMIS and both based on TE analysis, ARTEMIS is based on the genomic coverages of 1,280 TE types ^33^. Moreover, our TEANA models leveraged less than 100 cfDNA fragmentomic features, which is much less than previous models such as DELFI, ARTEMIS, and EXCEL, suggesting that focusing on genetic elements that are frequently dysregulated in cancer might benefit diagnosis. This discovery might serve as a valuable principle in optimizing cfDNA fragmentomics-based cancer diagnostic models, therefore deserves further investigations and validations in future studies.

In summary, in this study, we performed a comprehensive investigation on cfDNA fragmentomics in TEs. Our results suggested that chromatin states served as molecular regulators of cfDNA fragmentation, therefore extended our knowledge of cfDNA biology. In addition, in cancer patients, aberrant cfDNA fragmentomic features are more profound in TEs, demonstrating high translational significance in cancer diagnosis.

## Methods

### Ethics approval and blood sample processing

This study had been approved by the Ethics Committee of Shenzhen Bay Laboratory and Ethics Committee of The Third People’s Hospital of Shenzhen. Non-cancerous patients were recruited from The Third People’s Hospital of Shenzhen with written informed consents for cfChIP-seq experiments (N=6). For each subject, 10 mL peripheral blood was collected using EDTA-containing tubes and processed within 4 hours. Animal study was conducted according to protocols approved by SZBL Animal Center. Wildtype C57BL/6J strain mice (N=6) were housed under specific pathogen-free conditions with a 12 h light/dark cycle, at a temperature of 20-26 °C, and a relative humidity of 40-70%; mice were fed a standard mouse chow diet and were sacrificed at 12-30 weeks. For both human subjects and mice, blood samples were centrifuged at 1,600 g, 4 °C for 15 min, and then the plasma portion was harvested and re-recentrifuged at 16,000 g, 4 °C for 15 min to remove blood cells ^13,59^; plasma samples were stored at -80 °C until further usage.

### Murine cfDNA extraction and sequencing

CfDNA extraction and library preparation from murine plasma (N=6) were performed as in our previous study^46^. CfDNA was extracted with MagPure Circulating DNA LQ Kit (Magen, #IVD5432). Briefly, 40 μL proteinase K solution and 50 μL MagPure Particles G were added to 300-600 μL plasma; then 1.1 mL Lysis buffer was added to the mixture and incubated at room temperature for 15 min. After separation on the Magnetic Separation Rack, the supernatant was discarded and the magnetic beads were washed with 700 μL Buffers twice, and cfDNA was then eluted with 20 μL water. For each sample, 3-6 ng cfDNA was used to construct libraries using VAHTS Universal DNA Library Prep Kit for MGI (Vazyme, #NDM607) following the manufacturer’s instructions: 50 μL sample was incubated with 15 μL End Prep Mix 4 at 20 °C for 15 min and followed by 65 °C for 15 min; then DNA was ligated with sequencing adaptors at 20 °C for 15 min and purified with 60 μL Purification Beads; purified DNA was amplified for 12 cycles with PCR Mix and purified with 1x volume of magnetic beads. The PCR product was denatured at 98 °C for 3 min and circulated with DNA Rapid Ligase at 37 °C for 15 min (Vazyme, #NM201). Unused ligation primers were digested with Digestion Enzyme at 37 °C for 10 min. The cfDNA libraries were subjected to DNA nanoball generation and sequenced on an MGISEQ-T7 (MGI) sequencer in paired-end 100 bp mode.

### CfChIP-seq experiments and sequencing

CfChIP-seq experiments targeting H3K4me1, H3K9me3, and H3K27me3 were performed on human plasma samples, each with 2 biological replicates, following the protocol in Baca et al. ^39^ with minor modifications. Briefly, 20 μL Protein A and 20 μL Protein G magnetic beads were used for pre-clear with 1.8-2 mL plasma for 2 h at 4 °C. 4 μg antibody (H3K4me1, Abcam #ab8895; H3K9me3, Abcam #ab8898; H3K27me3, Active motif #39055) were coupled with 10 μL Protein A (Invitrogen, #10002D) and 10 μL Protein G (Invitrogen, #10004D) magnetic beads for 6 h at 4 °C with rotation in 0.5% BSA (Sigma, #A7906) in PBS. Then, the pre-cleared and conditioned plasma was subjected to antibody-coupled magnetic beads overnight with rotation at 4 °C. The reclaimed magnetic beads were washed with 1 mL of low salt washing buffer twice, 1 mL of high salt washing buffer twice, 1 mL of LiCl washing buffer twice, and 150 μL of 10 mM cold Tris-HCl twice. Subsequently, the beads were resuspended and incubated in 68 μL elution buffer for 1.5 h at 60 °C. The elution cfDNA was purified through MagPure Circulating DNA Mini Kit (Magen, #IVD5432), and libraries were prepared with ThruPLEX DNA-Seq Kit (Takara Bio, #R400674) following the manufacturer’s instructions. All cfChIP-seq libraries were sequenced on a NovaSeq X PLUS sequencer (Illumina) in paired-end 150 bp mode.

### Sequencing data analysis

For cfDNA whole genome sequencing, cfChIP-seq, and ATAC-seq experiments, data analyses (including preprocessing, alignment, and duplicate removal) were performed using a unified pipeline as in our previous studies ^13,60^. Briefly, raw reads were first processed using Ktrim software ^61^ (version 1.5.0) to remove adapters and low-quality cycles; the preprocessed reads were then aligned to the reference genome using Bowtie2 software ^62^ (version 2.3.5.1) in paired-end mode with default parameters; PCR duplicates were identified and removed using in-house programs ^60^, and the remaining reads were used for downstream analyses. EM-seq data was analyzed using Msuite2 software ^63,64^ (version 2.2.0), an all-in-one package that includes quality control, read alignment, and methylation calling. Prior to alignment, the tailing 25 bp in read 1 and leading 25 bp in read 2 were trimmed to avoid overhang issues inherent to cfDNA ^65^; hisat2 software ^66^ (version 2.2.1) was employed as the underlying aligner and all the other parameters were kept default. Only the samples with a mappability higher than 80%, a cytosine-to-thymine conversion rate higher than 98%, and at least 20 million usable reads were kept in this study (N=1201). The reference genomes NCBI GRCh38 (hg38), GRCm38 (mm10), and Dog10K_Boxer_Tasha (canFam6) were used for human, mouse, and dog, respectively. Note that only reads mapped to the autosomes were kept in the downstream analyses. For H3K27ac, H3K4me3, and H3K36me3 cfChIP-seq experiments, pre-aligned reads were downloaded from the literature ^38,39^; genomic coordinates were originally provided according to NCBI GRCh37 (hg19) reference genome and were translated to GRCh38 reference genome using the “liftOver” utility from UCSC genome browser ^67^.

### Transposon elements

For human, mouse, and dog, pre-annotated genomic coordinates of TEs for the corresponding reference genomes by RepeatMasker software ^68^ were downloaded from UCSC Genome Browser ^67^. The TE families were then ranked by their copy numbers. For human, we kept the TE families with more than 10,000 copies (N=16; Suppl. Table S1); for mouse and dog, the top-ranked 16 and 14 TE families were selected (Suppl. Table S1). Note that only TEs located in the autosomes were used in the downstream analyses.

### CfDNA fragmentomic features

For each cfDNA sample, we first picked up the reads overlapped with the 16 family of TEs separately using the “intersect” function in bedtools software ^69^ (version 2.29.2) with “-f 0.5” parameter, i.e., only the reads with at least 50% falling in the TEs were kept. Then, we focused on the following fragmentomic features for each family of TE using cfDNA reads falling into it: size, CCCA end motif usage, end motif diversity, E-index, and relative sequencing depth (RSD).

For each cfDNA read, the distance between the two outmost ends was determined as its size; reads shorter than or equal to 150 bp (i.e., ≤ 150 bp) were considered as “short fragments”. For each TE, the fraction of short fragments was utilized as the size parameter ^60^. For motif analysis, we extracted the 4-mer sequences (total 256 types) from the 5’-end of cfDNA reads, and for each 4-mer motif, we determined the fraction of reads with this motif; we then picked up the fraction of reads with 5’-CCCA sequence as CCCA end motif usage, and calculated the entropy of the 4-mer motifs as the motif diversity score ^3,15^.

For each TE, the E-index values were calculated as described in our previous study ^13^. Briefly, we analyzed cfDNA ends in an orientation-aware manner ^10^: for each cfDNA molecule, its fragment ends with lower and higher genome coordinates were termed as upstream (U) and downstream (D) ends, respectively, and processed separately in downstream analyses. We collected a pool of healthy controls (N=24) and recorded the counts of each genomic loci serving as U/D ends (the “model”); then for each sample to evaluate, we measured the consistency of its ends to the model using a weighted average approach (referred to as “E-index”) as illustrated in the following formula:

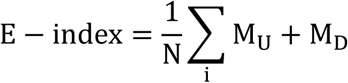

where *N* denoted the read number of the sample-of-interest, *i* denoted each sequencing read, *M_U_*, and *M_D_* denoted the counts of its ending positions serving as U and D ends in the model, respectively.

For each TE (or any element of interest), RSD was determined as the average sequencing depth for all its copies in relative to the genome-wide level. To calculate RSD, we counted the total number of cfDNA reads falling into the TE family, and then normalized it to the total size of all its copies and the total read number of the sample, as illustrated in the following formula:

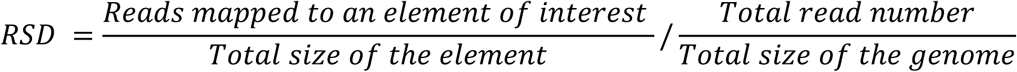

Based on this definition, an RSD value higher than 1 would suggest elevated depth than the genome-wide level, and an RSD value lower than 1 would suggest decreased depth than the genome-wide level.

### Association of cfDNA fragmentomics with epigenomic features

For DNA methylation analysis, we pooled the EM-seq data from 20 control samples with highest sequencing depth from Bie et al. dataset ^48^. The methylation densities of CpG sites were inferred, and for each TE family, we defined three subgroups based on the average methylation density of covered CpG sites: low (<20%), partial (20%-90%), and high (>90%). For histone modification data, sequencing reads from different cfChIP-seq experiments of the same type were pooled together. For open chromatin regions, we utilized ATAC-seq data from three major hematopoietic lineages that contribute to cfDNA: megakaryocytes, T-cells, and neutrophils ^3,38,40^. For each epigenomic feature, we categorized the copies in each TE family into two subgroups (i.e., low and high) based on the signal density of the corresponding epigenomic data in the copies. Then, to investigate the relationships between cfDNA fragmentomics and these epigenomic features, we analyzed epigenomics data in TEs in a paired manner (e.g., H3K27ac - SINE/ALU): we extracted the cfDNA reads overlapping each TE subgroups using data from 24 control subjects ^13^, and then computed the cfDNA fragmentomic features and compared these features across the subgroups.

### Chromatin state analysis

The chromatin states were defined using ChromHMM software ^43^ (version 1.25) using the cfChIP-seq data of 6 histone modifications (H3K27ac, H3K4me1, H3K4me3, H3K36me3, H3K27me3, and H3K9me3) versus cfDNA whole genome sequencing data from controls (as input signals). During the analysis, the state number was set to 15, window size was set to 200 bp, and the rest parameters were kept default ^44^. The imputed chromatin states were provided in Suppl. Table S3. Chromatin states were manually annotated based on the epigenomic signals ^44^. Then for each control sample, we extracted the cfDNA reads falling into each chromatin state (using the “intersect” function in bedtools software with “-f 0.5” parameter) then computed the cfDNA fragmentomic features for each chromatin state separately.

### AI-enabled cancer diagnosis and tumor origin tracing models

To build TEANA-Dx models for Cristiano et al. dataset, the following 5 cfDNA fragmentomic features were leveraged: fraction of short fragments, CCCA end motif usage, end motif diversity, RSD, and E-index values. For each sample, we calculated cfDNA fragmentomic features for 16 types of TEs (N=80 features in total). For Bie et al. dataset, besides those used in Cristiano et al. dataset, DNA methylation densities were also included, resulting in a total of 96 features. A gradient boosted decision tree (GBDT) algorithm, implemented in the “gbm” package (version 2.1.9) in R software (version 4.2.0), was used to build diagnostic models using the features. Hyperparameter tuning was performed using the “caret” package (version 6.0.94), as described in previous studies ^34,46^. A 10-fold nested cross-validation was applied as follows: the samples were randomly split into 10 equal-sized subsets, with 9 subsets used for training and the remaining subset used for testing. This process was repeated for each subset, and the prediction results for each testing subset were collected. Hence, for each sample, it was predicted by a model trained using samples without itself to eliminate information disclosure. The whole procedure was repeated 10 times, and the average prediction score for each sample was calculated and determined as its prediction score. The area under the receiver operating characteristic (ROC) curve (AUC) was used to quantify model performance. In addition, for Bie et al. ^48^ dataset, the samples were divided into non-overlapping training and testing subsets in the original study; we re-trained a TEANA-Dx model only based on the samples in the training subset using the same approach as described above, the trained model was then directly applied to the testing subset to evaluate its performance (including ROCs and sensitivities).

To build the tissue-origin prediction model (TEANA-Top) for Cristiano et al. dataset, the cancer samples were picked up and analyzed; for each cancer type, we treated the samples of this type as “positive” set while the rest samples of different cancer types as “negative” set to build classification model using the same approach as in TEANA-Dx described above. This process resulted in seven models, each for one cancer type. Then for each sample, we applied the 7 models and collected the prediction scores from all models, and the cancer type whose corresponding model showed the highest prediction score was reported as the tissue-origin prediction result.

## Data availability

Public cfDNA sequencing and cfChIP-seq datasets were obtained from CNGB Nucleotide Sequence Archive (accession number CNP0000680), GSA-Human (National Genomics Data Center, accession numbers HRA005521 and HRA003209), GEO (accession numbers GSE243474), Sequence Read Archive (accession number PRJNA823593), Zenodo (https://doi.org/10.5281/zenodo.3967253), and FinaleDB ^70^. Public ATAC-seq datasets were collected from GEO (accession numbers GSE76006 and GSE207654) and ENCODE Project (accession numbers ENCSR102RXG, ENCSR258RSH, ENCSR299LSN, and ENCSR934AWQ).

## Acknowledgements

This work was supported by National Natural Science Foundation of China (32401206), Guangdong Basic and Applied Basic Research Foundation (2023B1515120073), Shenzhen Clinical Research Center for Oral Diseases (20210617170745001), National Key R&D Program of China (2022YFA0912700), and Major Program of Shenzhen Bay Laboratory (S241101004), and Shenzhen Bay Scholar Fellowship (to C.D. and K.S.). We’d like to thank Ms. Qi Wang for technical assistance, and Shenzhen Bay Laboratory HPC facility for computational support.

## Declaration of interests

X.S. and K.S. had filed patent applications based on the method developed in this work; the remaining authors declare no conflict of interests.

## Author contributions

Conception and design: K.S.; Study supervision: X.S., K.S.; Development of methodology: F.G., Y.P., H.L., K.S.; Acquisition of data: all authors; Analysis and interpretation of data: all authors; Writing the manuscript: F.G., Y.P., H.L., K.S.; Review and edit of the manuscript: Y.Z., C.D., X.S., K.S.

