## Supplementary figures for "Cell-free DNA fragmentomic characteristics in transposon elements inform molecular regulators and enhance cancer diagnosis"

Gong *et al.*

This supplementary file covers Figures S1-S27.

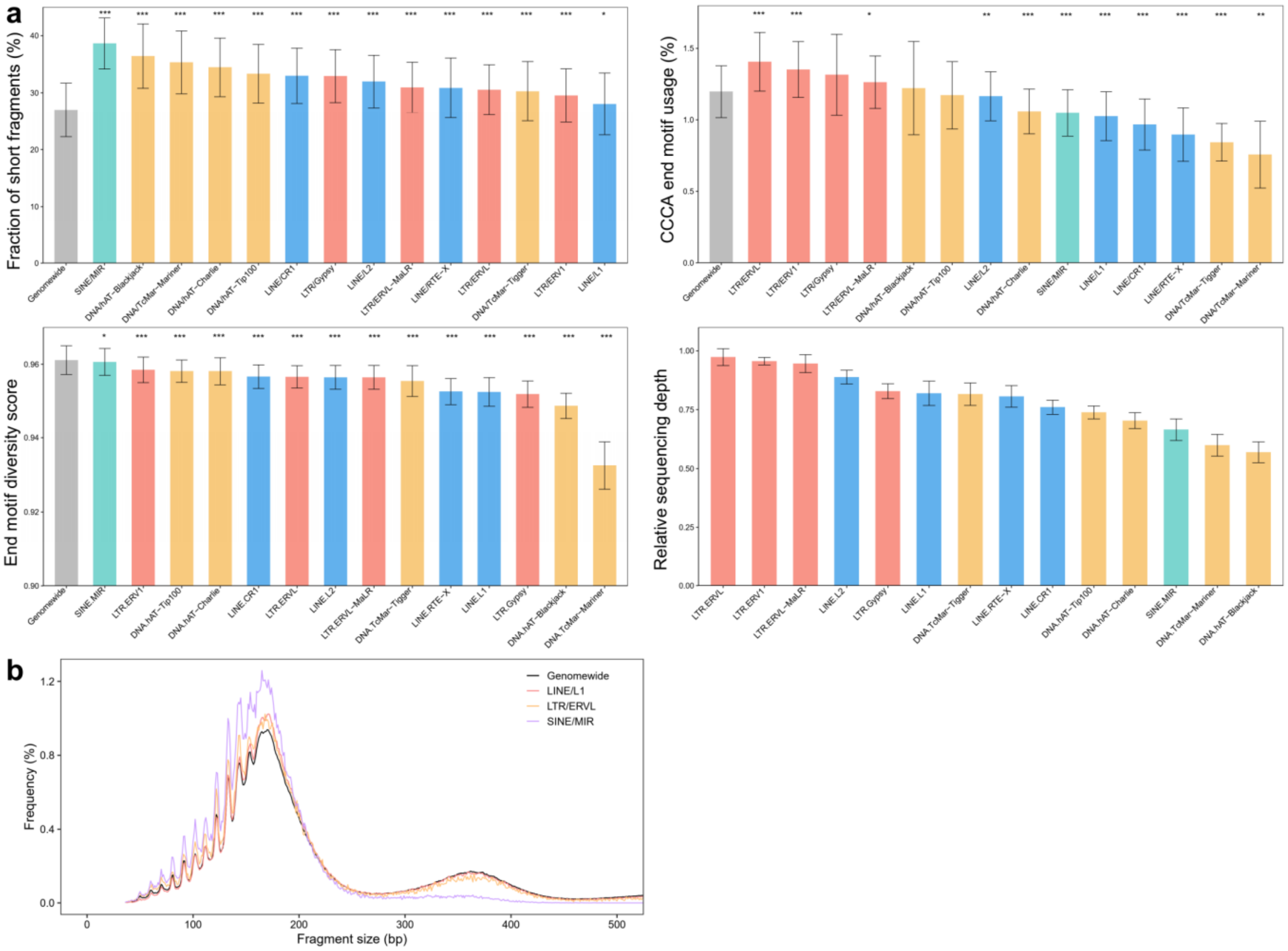


**Fig. S1. CfDNA fragmentomic characteristics in transposon elements (TEs) in dog samples (N=9).** (a) CfDNA size, motif patterns, and relative sequencing depths across 14 TEs. (b) Size distributions of cfDNA in representative transposon elements. In (a), bars indicated mean ± s.d., and p-values were calculated between each TE and genomewide level using paired t-tests. *$p<0.05$; **$p<0.01$, ***$p<0.001$.

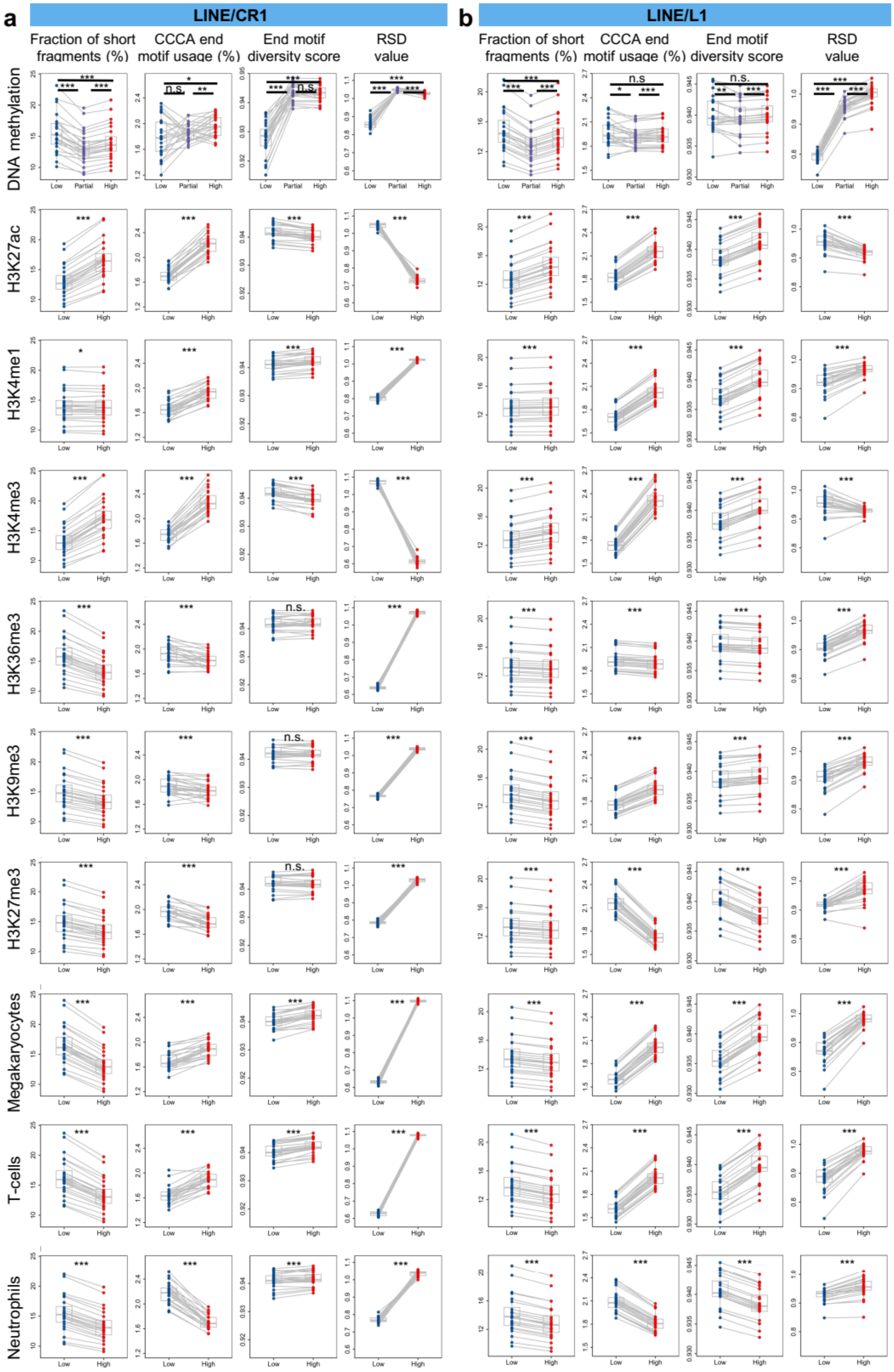
a
LINE/CR1
b
LINE/L1
Fraction of short fragments (%)
CCCA end motif usage (%)
End motif diversity score
RSD value
DNA methylation
H3K27ac
H3K4me1
H3K4me3
H3K36me3
H3K9me3
H3K27me3
Megakaryocytes
T-cells
Neutrophils
Low
Partial
High
n.s.

**Fig. S2. CfDNA fragmentomic characteristics in (a) LINE/CR1 and (b) LINE/L1 in different epigenomic context.** For DNA methylation analysis, copies in LINE/CR1 and LINE/L1 were divided into three groups based on the average DNA methylation levels of CpG sites they covered; for histone modification (H3K27ac, H3K4me1, H3K4me3, H3K36me3, H3K9me3, and H3K27me3) and open chromatin (Megakaryocyte, T-cells, and Neutrophils) analysis, copies in LINE/CR1 and LINE/L1 were divided into two groups based on the size-normalized signals of these epigenomic markers. P-values were computed using paired t-tests. *$p<0.05$, **$p<0.01$, ***$p<0.001$.

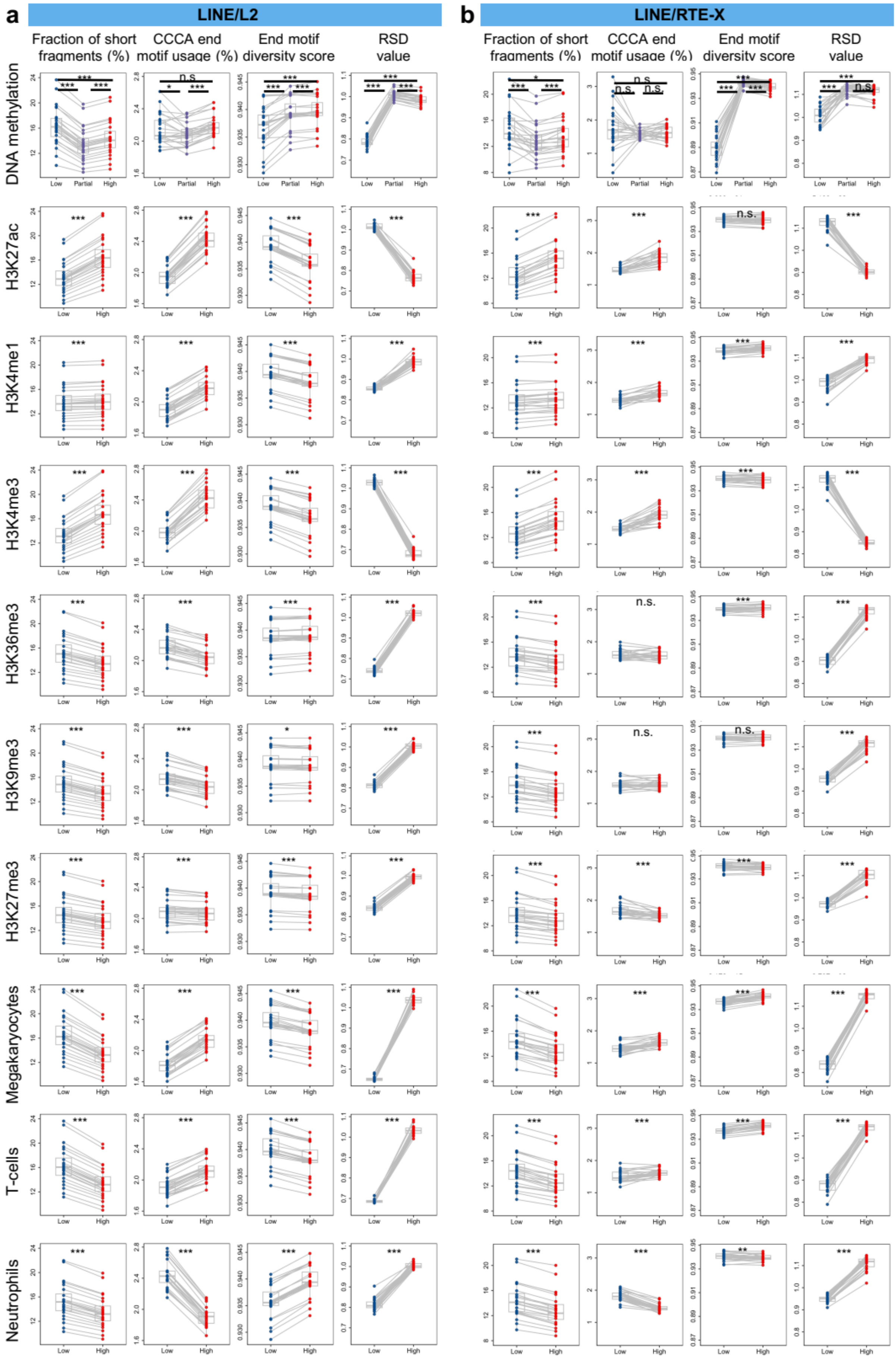

a
LINE/L2
b
LINE/RTE-X
Fraction of short fragments (%)
CCCA end motif usage (%)
End motif diversity score
RSD value
DNA methylation
H3K27ac
H3K4me1
H3K4me3
H3K36me3
H3K9me3
H3K27me3
Megakaryocytes
T-cells
Neutrophils
Low
Partial
High
n.s.

**Fig. S3. CfDNA fragmentomic characteristics in (a) LINE/L2 and (b) LINE/RTE-X in different epigenomic context.** For DNA methylation analysis, copies in LINE/L2 and LINE/RTE-X were divided into three groups based on the average DNA methylation levels of CpG sites they covered; for histone modification (H3K27ac, H3K4me1, H3K4me3, H3K36me3, H3K9me3, and H3K27me3) and open chromatin (Megakaryocyte, T-cells, and Neutrophils) analysis, copies in LINE/L2 and LINE/RTE-X were divided into two groups based on the size-normalized signals of these epigenomic markers. P-values were computed using paired t-tests. *$p<0.05$, **$p<0.01$, ***$p<0.001$.

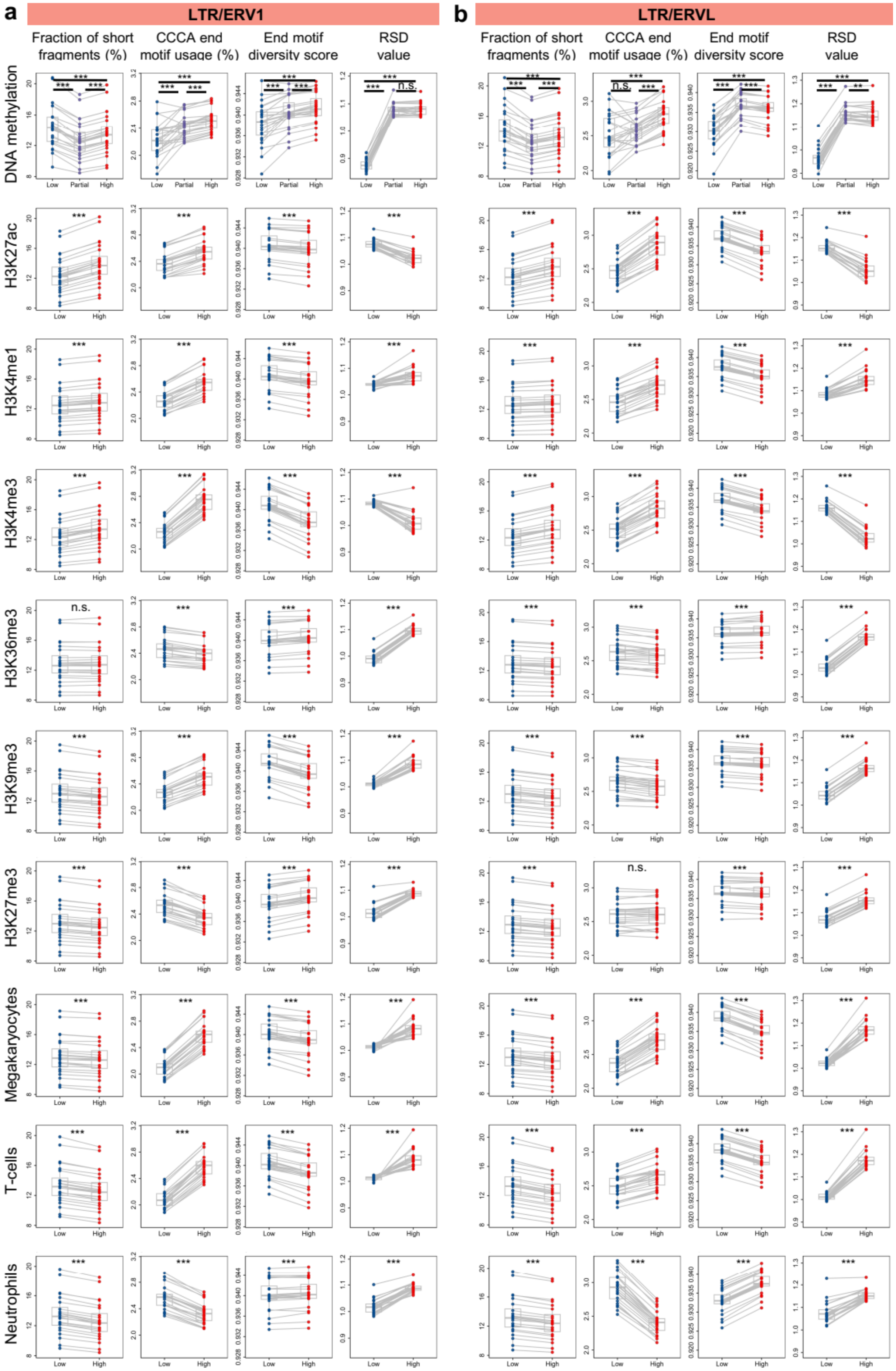
a
LTR/ERV1
b
LTR/ERVL
Fraction of short fragments (%)
CCCA end motif usage (%)
End motif diversity score
RSD value
DNA methylation
H3K27ac
H3K4me1
H3K4me3
H3K36me3
H3K9me3
H3K27me3
Megakaryocytes
T-cells
Neutrophils
Low
Partial
High
***
**
n.s.

**Fig. S4. CfDNA fragmentomic characteristics in (a) LTR/ERV1 and (b) LTR/ERVL in different epigenomic context.** For DNA methylation analysis, copies in LTR/ERV1 and LTR/ERVL were divided into three groups based on the average DNA methylation levels of CpG sites they covered; for histone modification (H3K27ac, H3K4me1, H3K4me3, H3K36me3, H3K9me3, and H3K27me3) and open chromatin (Megakaryocyte, T-cells, and Neutrophils) analysis, copies in LTR/ERV1 and LTR/ERVL were divided into two groups based on the size-normalized signals of these epigenomic markers. P-values were computed using paired t-tests. *$p<0.05$, **$p<0.01$, ***$p<0.001$.

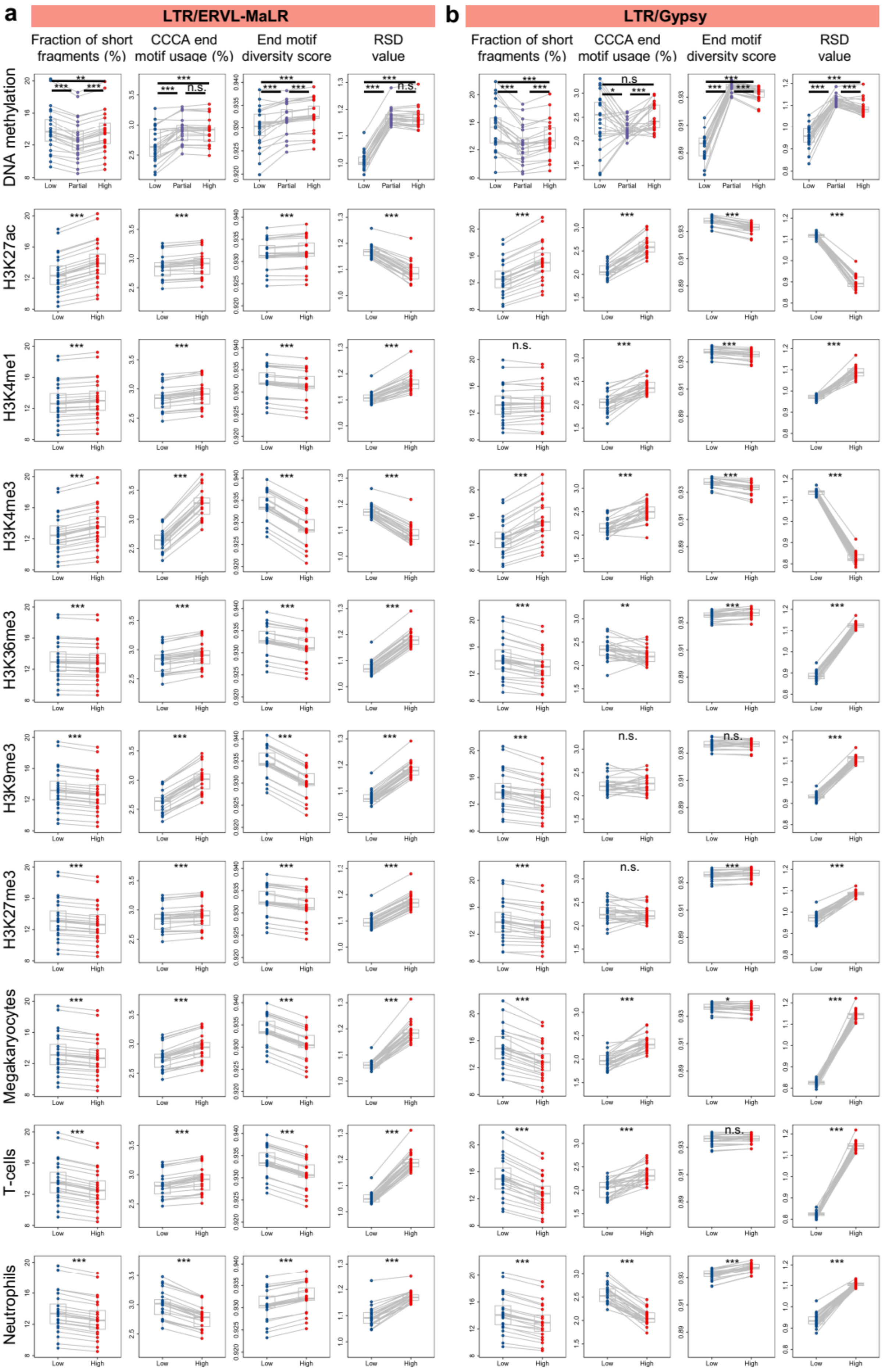
a
LTR/ERVL-MaLR
b
LTR/Gypsy
Fraction of short fragments (%)
CCCA end motif usage (%)
End motif diversity score
RSD value
DNA methylation
H3K27ac
H3K4me1
H3K4me3
H3K36me3
H3K9me3
H3K27me3
Megakaryocytes
T-cells
Neutrophils
Low
Partial
High

**Fig. S5. CfDNA fragmentomic characteristics in (a) LTR/ERVL-MaLR and (b) LTR/Gypsy in different epigenomic context.** For DNA methylation analysis, copies in LTR/ERVL-MaLR and LTR/Gypsy were divided into three groups based on the average DNA methylation levels of CpG sites they covered; for histone modification (H3K27ac, H3K4me1, H3K4me3, H3K36me3, H3K9me3, and H3K27me3) and open chromatin (Megakaryocyte, T-cells, and Neutrophils) analysis, copies in LTR/ERVL-MaLR and LTR/Gypsy were divided into two groups based on the size-normalized signals of these epigenomic markers. P-values were computed using paired t-tests. *p<0.05, **p<0.01, ***p<0.001.

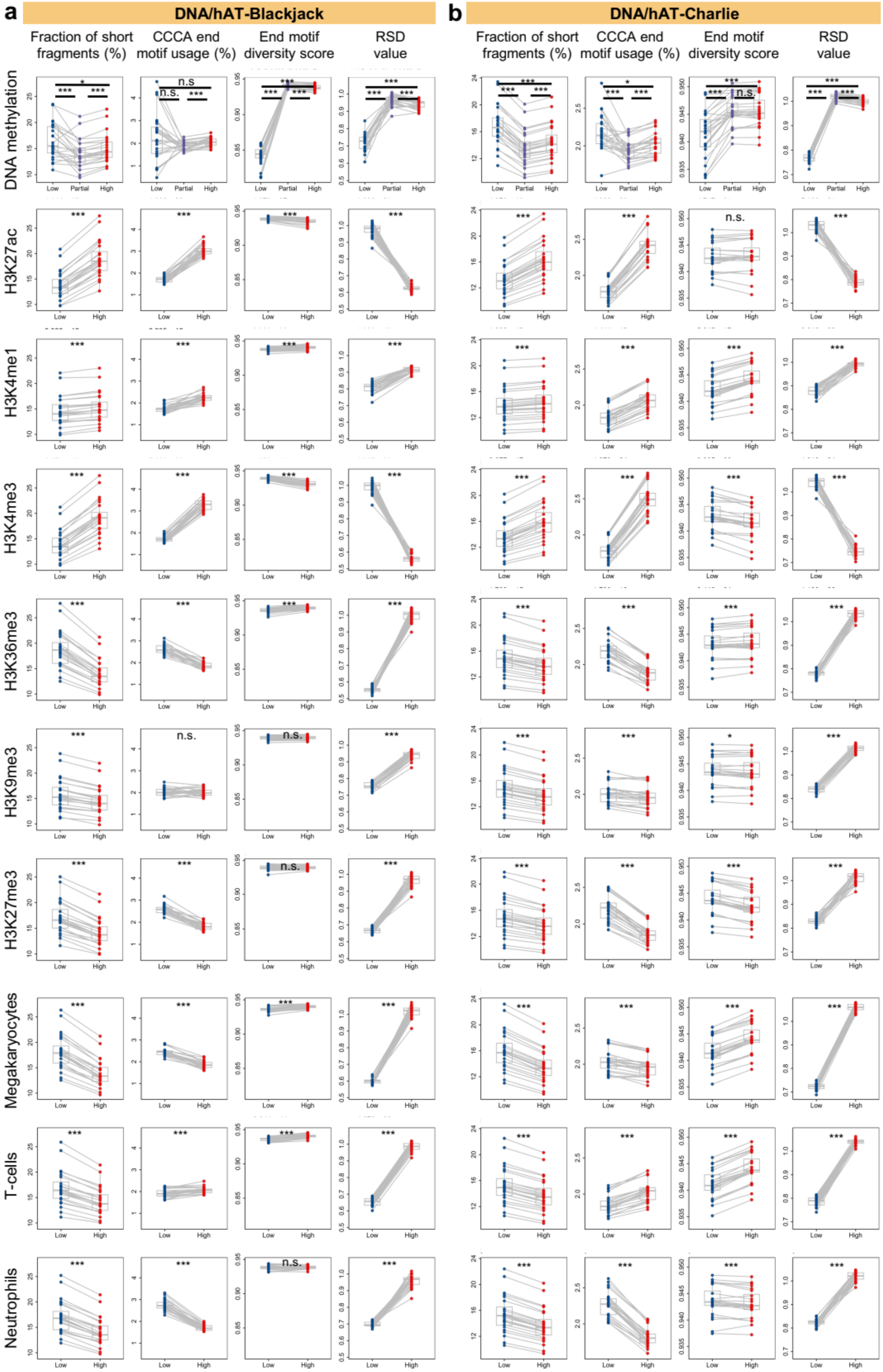
a
DNA/hAT-Blackjack
b
DNA/hAT-Charlie
Fraction of short fragments (%)
CCCA end motif usage (%)
End motif diversity score
RSD value
DNA methylation
H3K27ac
H3K4me1
H3K4me3
H3K36me3
H3K9me3
H3K27me3
Megakaryocytes
T-cells
Neutrophils
Low
Partial
High
n.s.

**Fig. S6. CfDNA fragmentomic characteristics in (a) DNA/hAT-Blackjack and (b) DNA/hAT-Charlie in different epigenomic context.** For DNA methylation analysis, copies in DNA/hAT-Blackjack and DNA/hAT-Charlie were divided into three groups based on the average DNA methylation levels of CpG sites they covered; for histone modification (H3K27ac, H3K4me1, H3K4me3, H3K36me3, H3K9me3, and H3K27me3) and open chromatin (Megakaryocyte, T-cells, and Neutrophils) analysis, copies in DNA/hAT-Blackjack and DNA/hAT-Charlie were divided into two groups based on the size-normalized signals of these epigenomic markers. P-values were computed using paired t-tests. *$p<0.05$, **$p<0.01$, ***$p<0.001$.

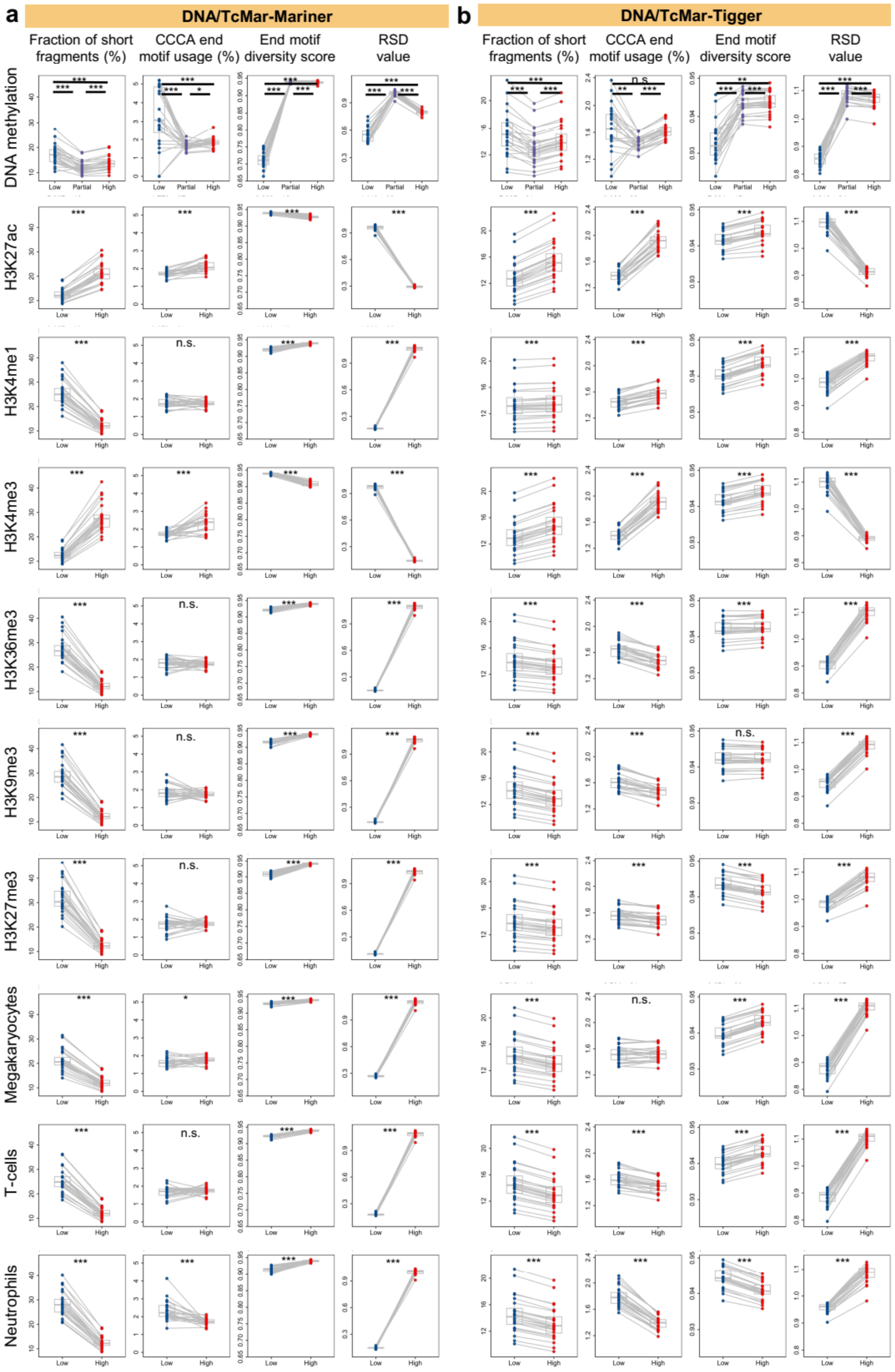
a
DNA/TcMar-Mariner
b
DNA/TcMar-Tigger
Fraction of short fragments (%)
CCCA end motif usage (%)
End motif diversity score
RSD value
DNA methylation
H3K27ac
H3K4me1
H3K4me3
H3K36me3
H3K9me3
H3K27me3
Megakaryocytes
T-cells
Neutrophils
Low
Partial
High
***
**
*
n.s.

**Fig. S7. CfDNA fragmentomic characteristics in (a) DNA/TcMar-Mariner and (b) DNA/TcMar-Tigger in different epigenomic context.** For DNA methylation analysis, copies in DNA/TcMar-Mariner and DNA/TcMar-Tigger were divided into three groups based on the average DNA methylation levels of CpG sites they covered; for histone modification (H3K27ac, H3K4me1, H3K4me3, H3K36me3, H3K9me3, and H3K27me3) and open chromatin (Megakaryocyte, T-cells, and Neutrophils) analysis, copies in DNA/TcMar-Mariner and DNA/TcMar-Tigger were divided into two groups based on the size-normalized signals of these epigenomic markers. P-values were computed using paired t-tests. *$p<0.05$, **$p<0.01$, ***$p<0.001$.

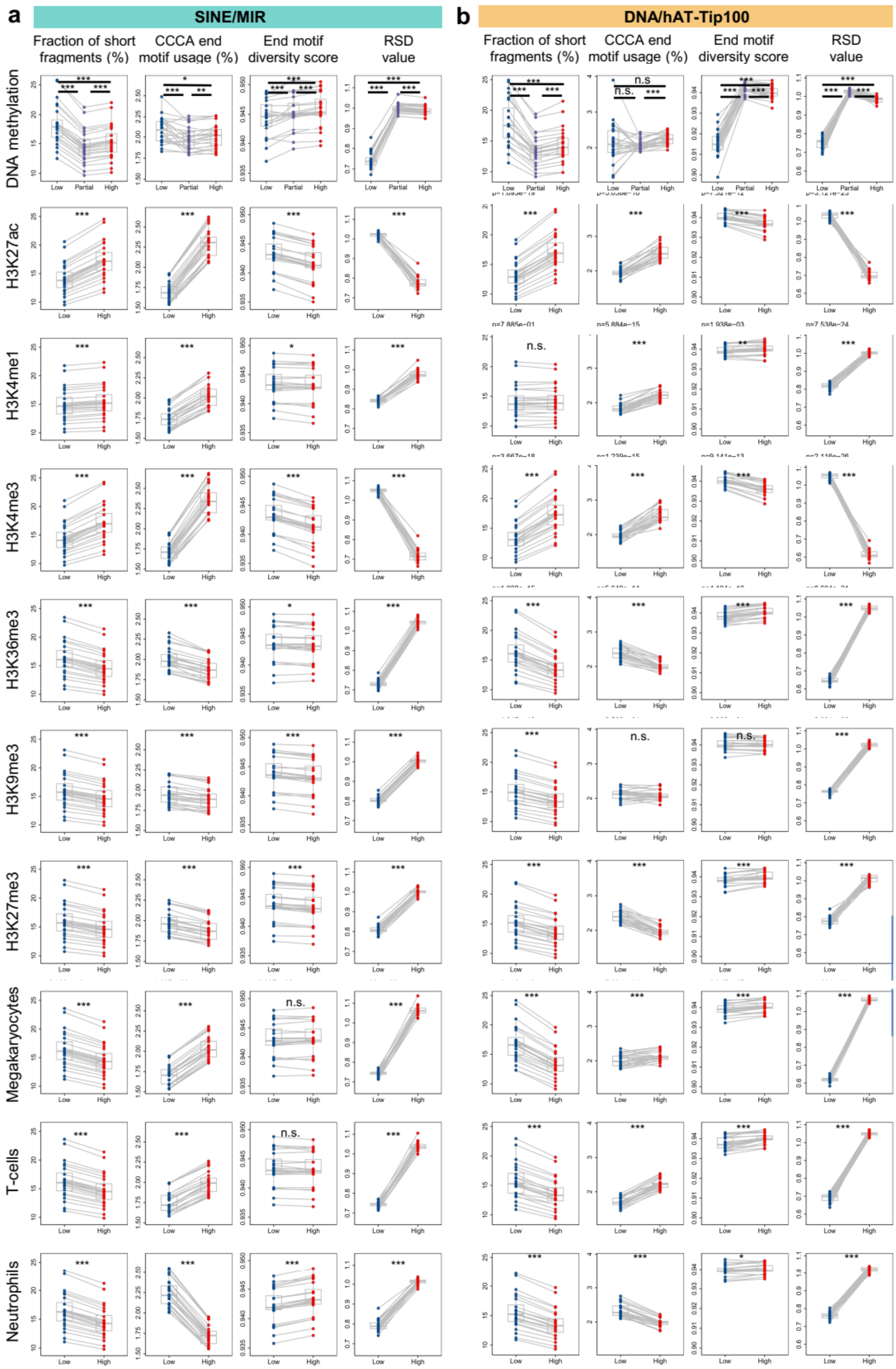
a
SINE/MIR
b
DNA/hAT-Tip100
Fraction of short fragments (%)
CCCA end motif usage (%)
End motif diversity score
RSD value
DNA methylation
H3K27ac
H3K4me1
H3K4me3
H3K36me3
H3K9me3
H3K27me3
Megakaryocytes
T-cells
Neutrophils
Low
Partial
High

**Fig. S8. CfDNA fragmentomic characteristics in (a) SINE/MIR and (b) DNA/hAT-Tip100 in different epigenomic context.** For DNA methylation analysis, copies in SINE/MIR and DNA/hAT-Tip100 were divided into three groups based on the average DNA methylation levels of CpG sites they covered; for histone modification (H3K27ac, H3K4me1, H3K4me3, H3K36me3, H3K9me3, and H3K27me3) and open chromatin (Megakaryocyte, T-cells, and Neutrophils) analysis, copies in SINE/MIR and DNA/hAT-Tip100were divided into two groups based on the size-normalized signals of these epigenomic markers. P-values were computed using paired t-tests. *p<0.05, **p<0.01, ***p<0.001.

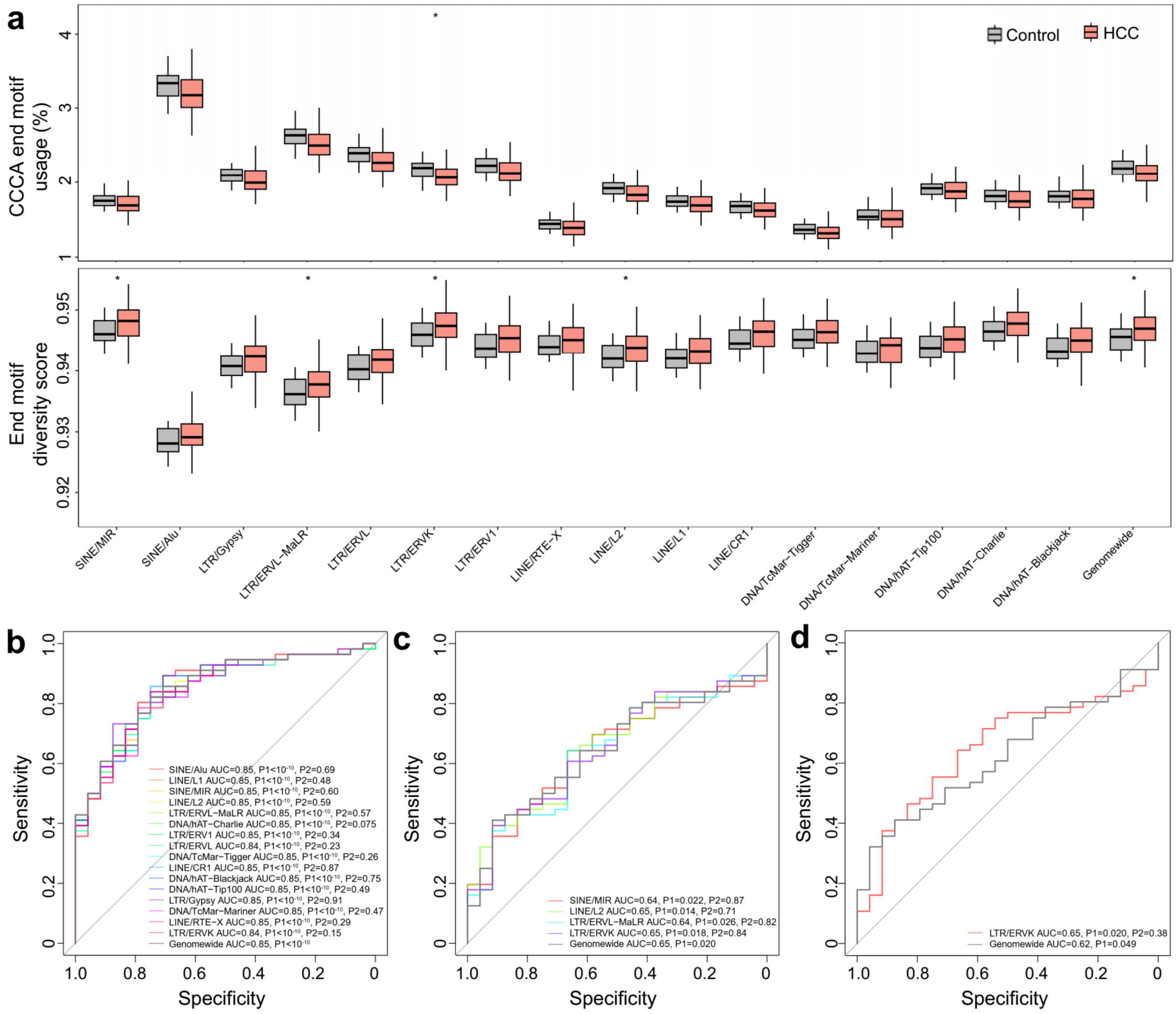


**Fig. S9. CfDNA fragmentomic characteristics in TEs in hepatocellular carcinoma (HCC) patients.** (a) motif patterns between controls and HCC patients across various TEs. Grey and orange boxes presented controls and HCC samples, respectively. Receiver Operating Characteristic (ROC) curves for fraction of short cfDNA fragments (b), end motif diversity score (c) and CCCA end motif usages (d) in various TEs and genomewide to differentiate HCC samples from controls. In (a), p-values were calculated using Mann-Whitney U tests. In (b-d), P1 were p-values for AUCs calculated using Z-tests, and P2 were p-values comparing the ROCs of each TE versus genomewide level using DeLong tests. *p<0.05, **p<0.01, ***p<0.001.

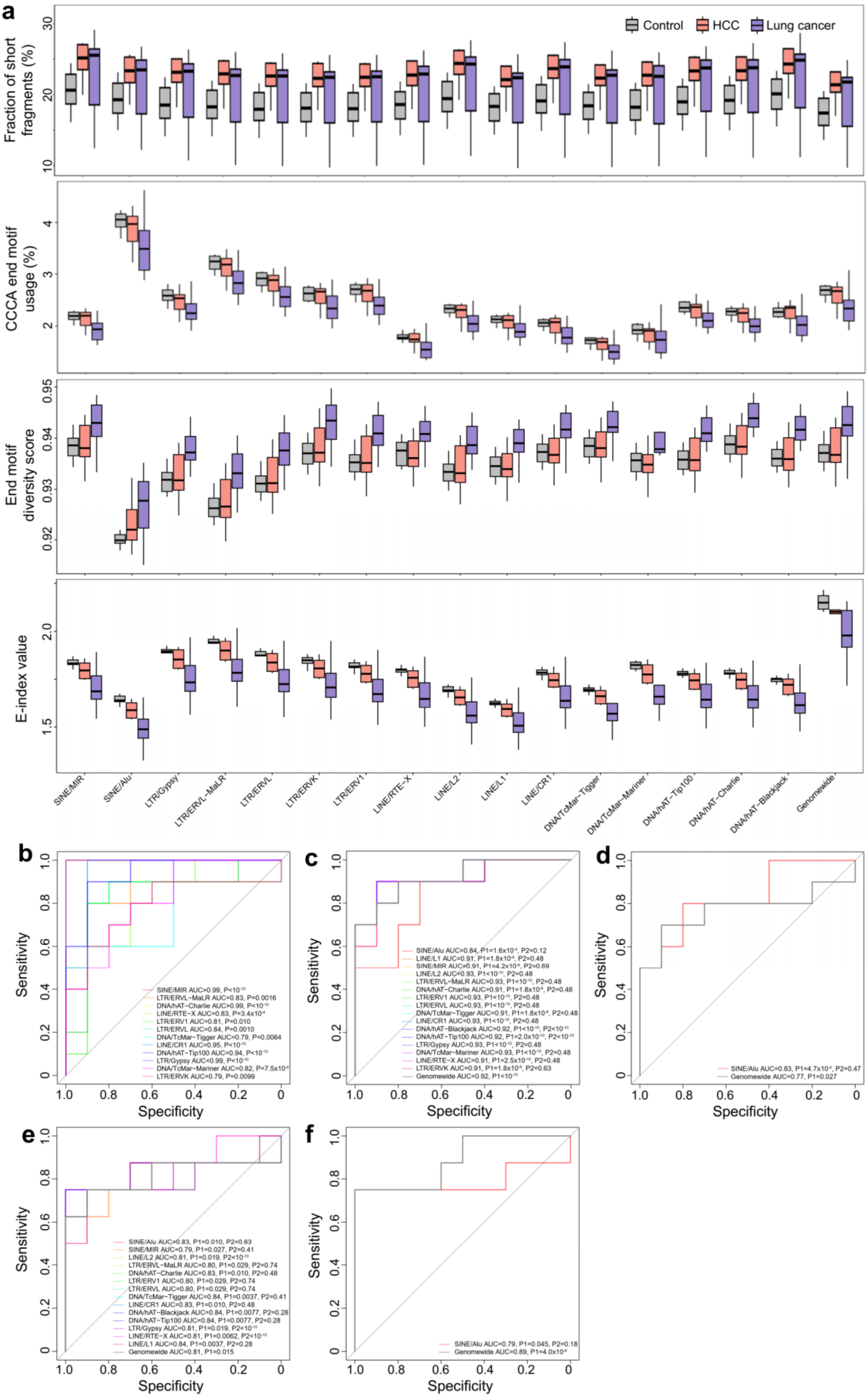
a
Control
HCC
Lung cancer
Fraction of short fragments (%)
CCCA end motif usage (%)
End motif diversity score
E-index value
SINE/MIR
SINE/Alu
LTR/Gypsy
LTR/ERVL−MaLR
LTR/ERVL
LTR/ERVK
LTR/ERV1
LINE/RTE−X
LINE/L2
LINE/L1
LINE/CR1
DNA/TcMar−Tigger
DNA/TcMar−Mariner
DNA/hAT−Tip100
DNA/hAT−Charlie
DNA/hAT−Blackjack
Genomewide
b
Sensitivity
Specificity
SINE/MIR AUC>0.99, P<$10^{-10}$
LTR/ERVL−MaLR AUC=0.83, P=0.0016
DNA/hAT−Charlie AUC>0.99, P<$10^{-10}$
LINE/RTE−X AUC=0.83, P=$3.4x10^{-4}$
LTR/ERV1 AUC=0.81, P=0.010
LTR/ERVL AUC=0.84, P=0.0010
DNA/TcMar−Tigger AUC=0.79, P=0.0064
LINE/CR1 AUC=0.95, P<$10^{-10}$
DNA/hAT−Tip100 AUC=0.94, P<$10^{-10}$
LTR/Gypsy AUC<0.99, P<$10^{-10}$
DNA/TcMar−Mariner AUC=0.82, P=$7.5x10^{-4}$
LTR/ERVK AUC=0.79, P=0.0099
c
Sensitivity
Specificity
SINE/Alu AUC=0.84, P1=$1.6x10^{-4}$, P2=0.12
LINE/L1 AUC=0.91, P1=$1.8x10^{-9}$, P2=0.48
SINE/MIR AUC=0.91, P1=$4.2x10^{-9}$, P2=0.69
LINE/L2 AUC=0.93, P1<$10^{-10}$, P2=0.48
LTR/ERVL−MaLR AUC=0.93, P1<$10^{-10}$, P2=0.48
DNA/hAT−Charlie AUC=0.91, P1=$1.8x10^{-9}$, P2=0.48
LTR/ERV1 AUC=0.93, P1<$10^{-10}$, P2=0.48
LTR/ERVL AUC=0.93, P1<$10^{-10}$, P2=0.48
DNA/TcMar−Tigger AUC=0.91, P1=$1.8x10^{-9}$, P2=0.48
LINE/CR1 AUC=0.93, P1<$10^{-10}$, P2=0.48
DNA/hAT−Blackjack AUC=0.92, P1<$10^{-10}$, P2<$10^{-10}$
DNA/hAT−Tip100 AUC=0.92, P1=$2.0x10^{-10}$, P2<$10^{-10}$
LTR/Gypsy AUC=0.93, P1<$10^{-10}$, P2=0.48
DNA/TcMar−Mariner AUC=0.93, P1<$10^{-10}$, P2=0.48
LINE/RTE−X AUC=0.91, P1=$2.5x10^{-10}$, P2=0.48
LTR/ERVK AUC=0.91, P1=$1.8x10^{-9}$, P2=0.63
Genomewide AUC=0.92, P1<$10^{-10}$
d
Sensitivity
Specificity
SINE/Alu AUC=0.83, P1=$4.7x10^{-4}$, P2=0.47
Genomewide AUC=0.77, P1=0.027
e
Sensitivity
Specificity
SINE/Alu AUC=0.83, P1=0.010, P2=0.63
SINE/MIR AUC=0.79, P1=0.027, P2=0.41
LINE/L2 AUC=0.81, P1=0.019, P2<$10^{-10}$
LTR/ERVL−MaLR AUC=0.80, P1=0.029, P2=0.74
DNA/hAT−Charlie AUC=0.83, P1=0.010, P2=0.48
LTR/ERV1 AUC=0.80, P1=0.029, P2=0.74
LTR/ERVL AUC=0.80, P1=0.029, P2=0.74
DNA/TcMar−Tigger AUC=0.84, P1=0.0037, P2=0.41
LINE/CR1 AUC=0.83, P1=0.010, P2=0.48
DNA/hAT−Blackjack AUC=0.84, P1=0.0077, P2=0.28
DNA/hAT−Tip100 AUC=0.84, P1=0.0077, P2=0.28
LTR/Gypsy AUC=0.81, P1=0.019, P2<$10^{-10}$
LINE/RTE−X AUC=0.81, P1=0.0062, P2<$10^{-10}$
LINE/L1 AUC=0.84, P1=0.0037, P2=0.28
Genomewide AUC=0.81, P1=0.015
f
Sensitivity
Specificity
SINE/Alu AUC=0.79, P1=0.045, P2=0.18
Genomewide AUC=0.89, P1=$4.0x10^{-6}$

**Fig. S10. CfDNA fragmentomic characteristics in TEs in Liang et al. dataset.** (a) Fraction of short cfDNA fragments (i.e., ≤ 150bp), motif patterns, and E-index values between controls and cancer patients across various TEs. Grey, orange, and purple boxes presented controls, HCC samples and lung cancer samples, respectively. Receiver Operating Characteristic (ROC) curves for RDS values (b) in various TEs to differentiate HCC samples from controls. Receiver Operating Characteristic (ROC) curves for fraction of short cfDNA fragments (c), and E-index values(d) in various TEs and genomewide to differentiate HCC samples from controls. Receiver Operating Characteristic (ROC) curves for end motif diversity score (e) and E-index values (f) in various TEs and genomewide to differentiate lung cancer samples from controls. In (b), p-values for Area Under the ROC Curves (AUCs) were calculated using Z-tests. In (c-f), P1 were p-values for AUCs calculated using Z-tests, and P2 were p-values comparing the ROCs of each TE versus genomewide level using DeLong tests. *p<0.05, **p<0.01, ***p<0.001.

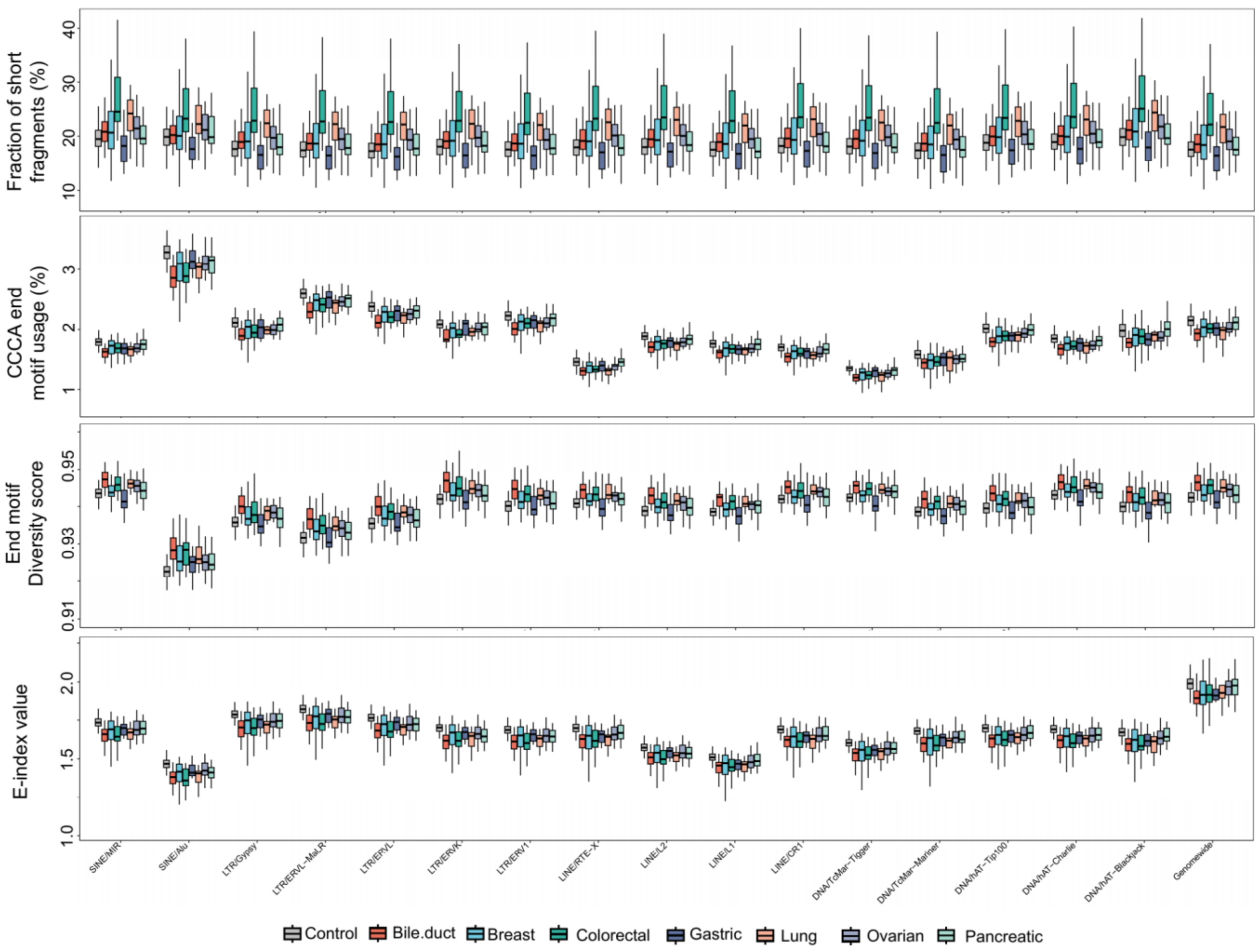


**Fig. S11. CfDNA fragmentomic characteristics in TEs in Cristiano et al. dataset.** Fraction of short cfDNA fragments (i.e., ≤ 150bp), motif patterns, and E-index values between controls and cancer patients across various TEs. Grey, red, blue, green, dark purple, orange, light purple and light green boxes presented controls, bile duct cancer, breast cancer, colorectal cancer, gastric cancer, lung cancer, ovarian cancer, and pancreatic cancer samples, respectively. Note that the abnormal size distribution of gastric cancer samples has been reported in previous study (Ju et al. Cell Reports Methods 2024).

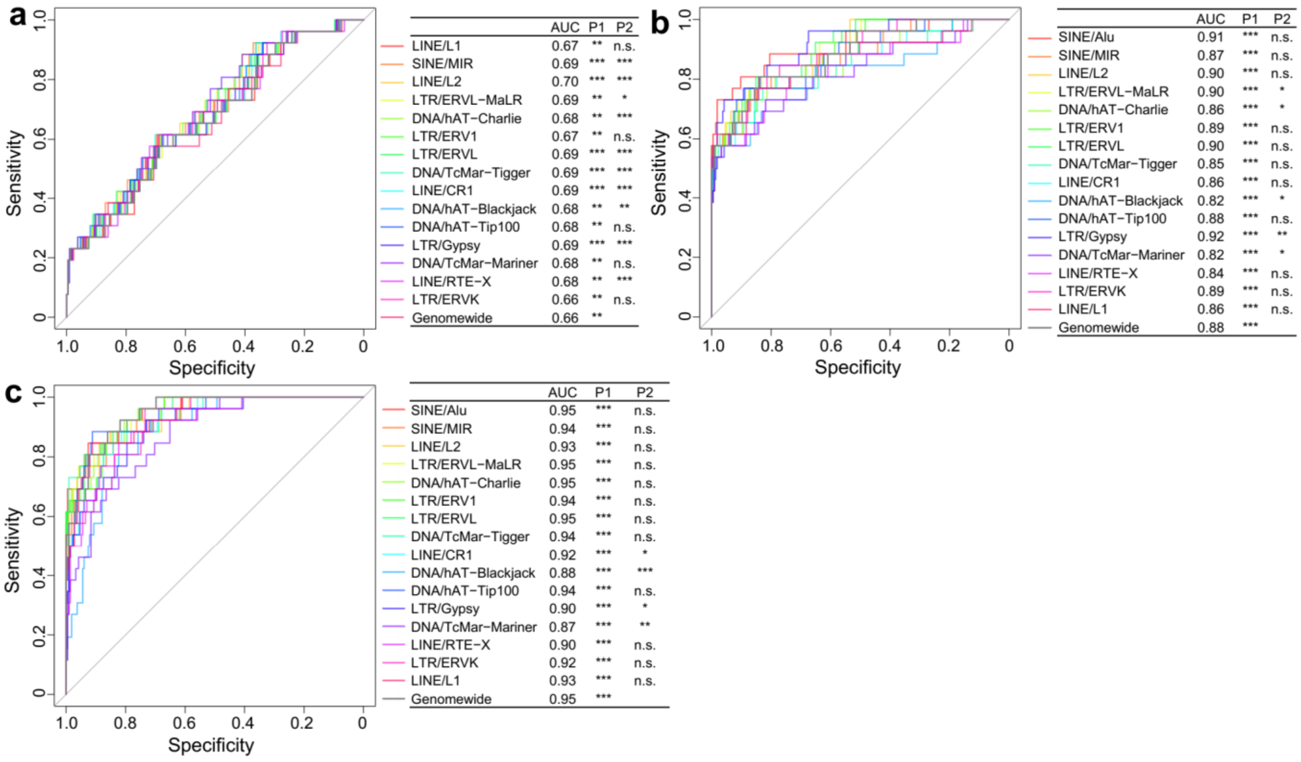


**Fig. S12. ROC curves for TEs to differentiate bile duct cancer samples from controls in Cristiano et al. dataset.** Receiver Operating Characteristic (ROC) curves for (a) short cfDNA fragments, (b) end motif diversity score, and (c) CCCA end motif usages in TEs and genomewide level to differentiate bile duct cancer samples from controls. P1 represented p-value for the AUC calculated using Z-tests, and P2 represented p-value comparing the ROCs of TE versus genomewide level using DeLong tests. *p<0.05, **p<0.01, ***p<0.001.

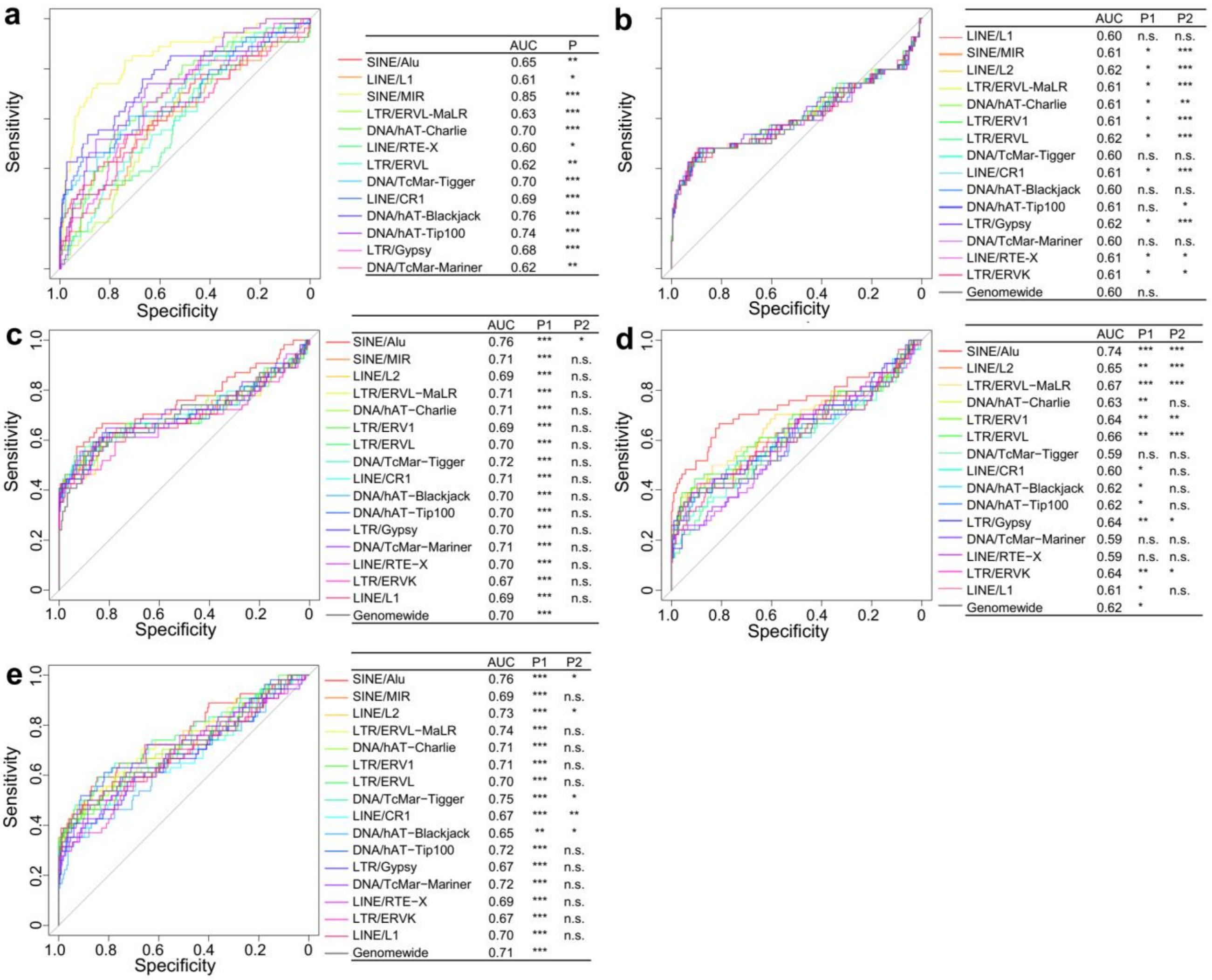


**Fig. S13. ROC curves for TEs to differentiate breast cancer samples from controls in Cristiano et al. dataset.** Receiver Operating Characteristic (ROC) curves for (a) RSD values, (b) E-index values, (c) short cfDNA fragments, (d) end motif diversity score, and (e) CCCA end motif usages in TEs and genomewide level to differentiate bile duct cancer samples from controls. P1 represented p-value for the AUC calculated using Z-tests, and P2 represented p-value comparing the ROCs of TE versus genomewide level using DeLong tests. *p<0.05, **p<0.01, ***p<0.001.

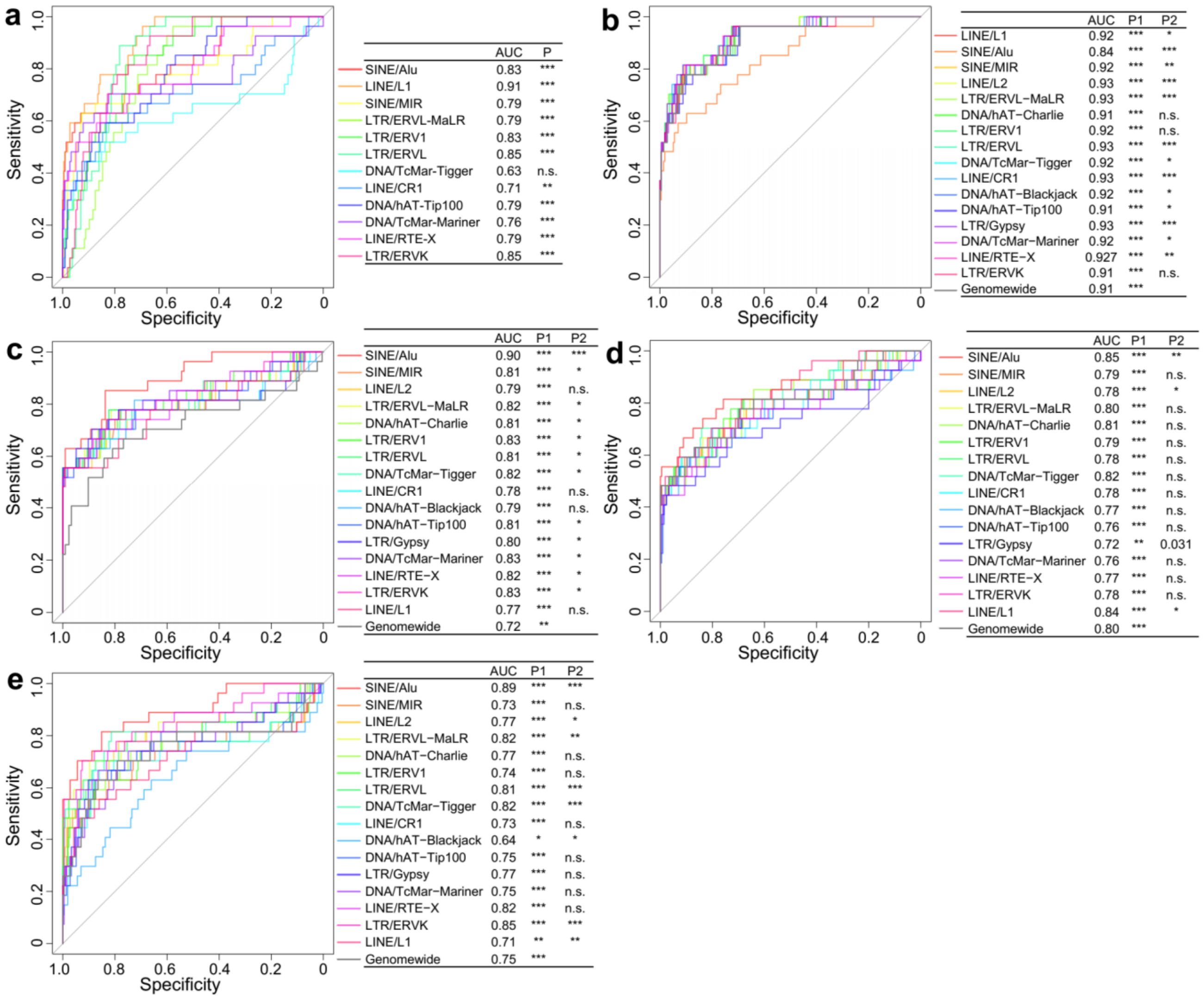


**Fig. S14. ROC curves for TEs to differentiate colorectal cancer samples from controls in Cristiano et al. dataset.** Receiver Operating Characteristic (ROC) curves for (a) RSD values, (b) E-index values, (c) short cfDNA fragments, (d) end motif diversity score, and (e) CCCA end motif usages in TEs and genomewide level to differentiate bile duct cancer samples from controls. P1 represented p-value for the AUC calculated using Z-tests, and P2 represented p-value comparing the ROCs of TE versus genomewide level using DeLong tests. *p<0.05, **p<0.01, ***p<0.001.

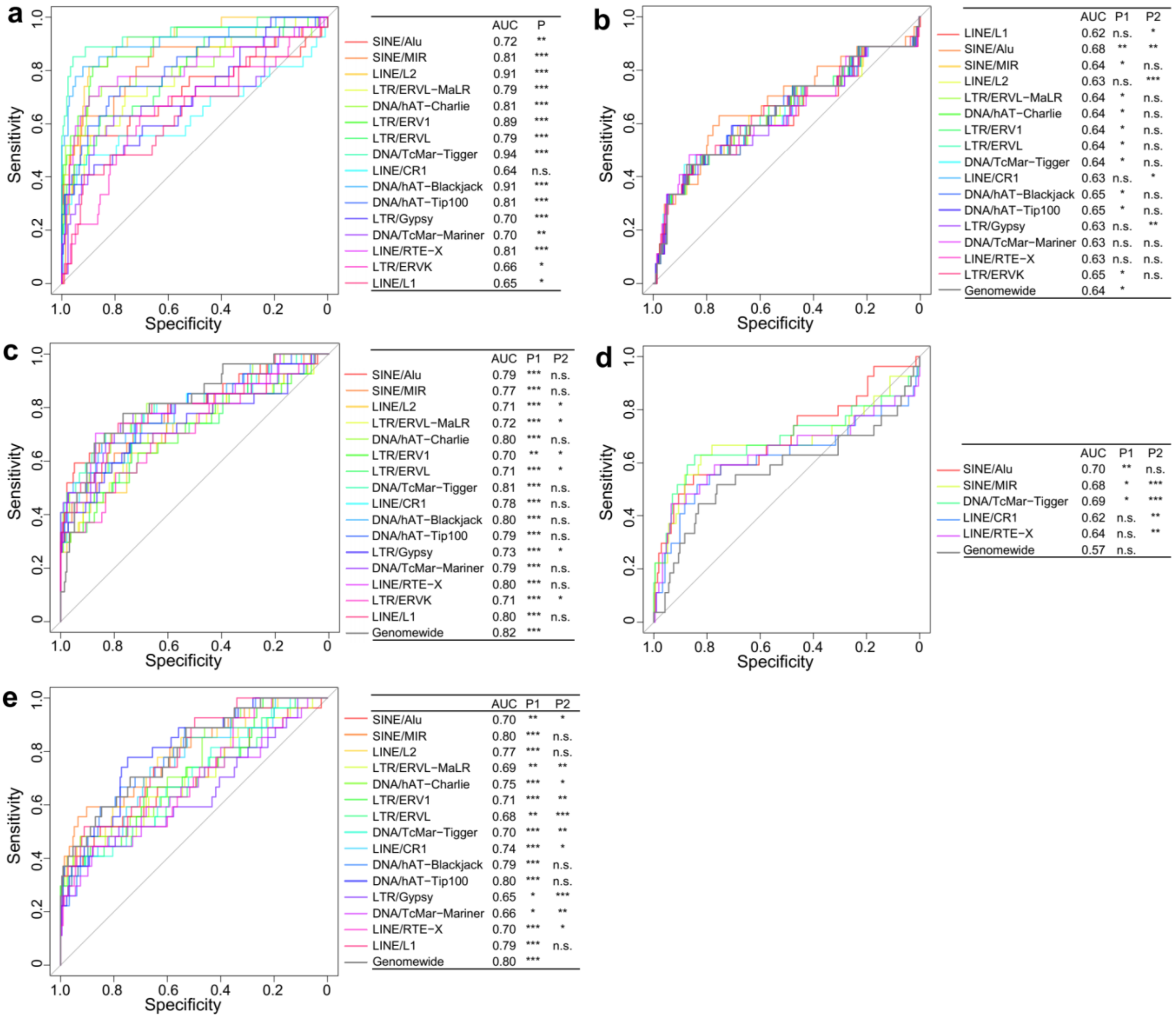


**Fig. S15. ROC curves for TEs to differentiate gastric cancer samples from controls in Cristiano et al. dataset.** Receiver Operating Characteristic (ROC) curves for (a) RSD values, (b) E-index values, (c) short cfDNA fragments, (d) end motif diversity score, and (e) CCCA end motif usages in TEs and genomewide level to differentiate bile duct cancer samples from controls. P1 represented p-value for the AUC calculated using Z-tests, and P2 represented p-value comparing the ROCs of TE versus genomewide level using DeLong tests. *p<0.05, **p<0.01, ***p<0.001.

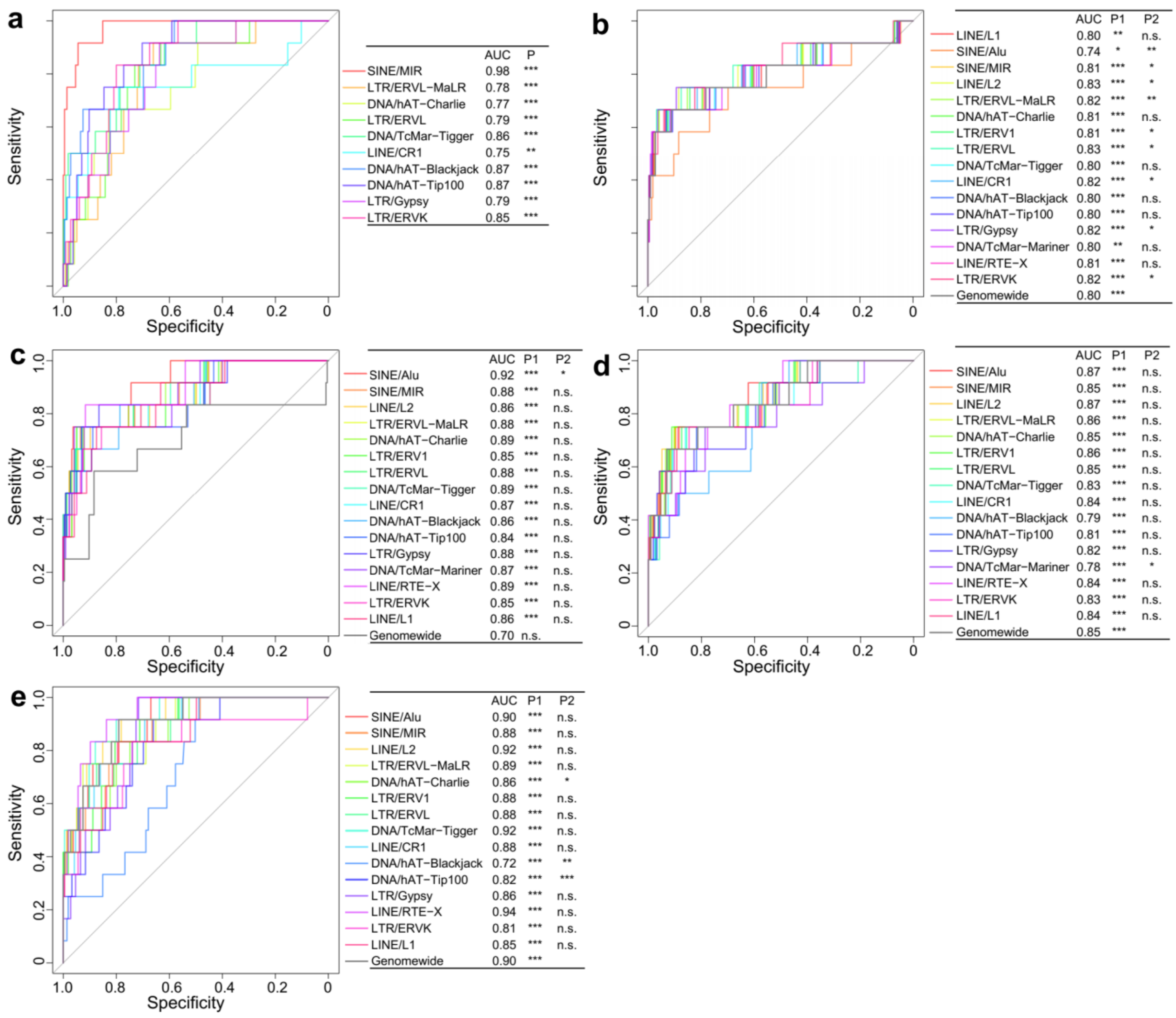


**Fig. S16. ROC curves for TEs to differentiate lung cancer samples from controls in Cristiano et al. dataset.** Receiver Operating Characteristic (ROC) curves for (a) RSD values, (b) E-index values, (c) short cfDNA fragments, (d) end motif diversity score, and (e) CCCA end motif usages in TEs and genomewide level to differentiate bile duct cancer samples from controls. P1 represented p-value for the AUC calculated using Z-tests, and P2 represented p-value comparing the ROCs of TE versus genomewide level using DeLong tests. *p<0.05, **p<0.01, ***p<0.001.

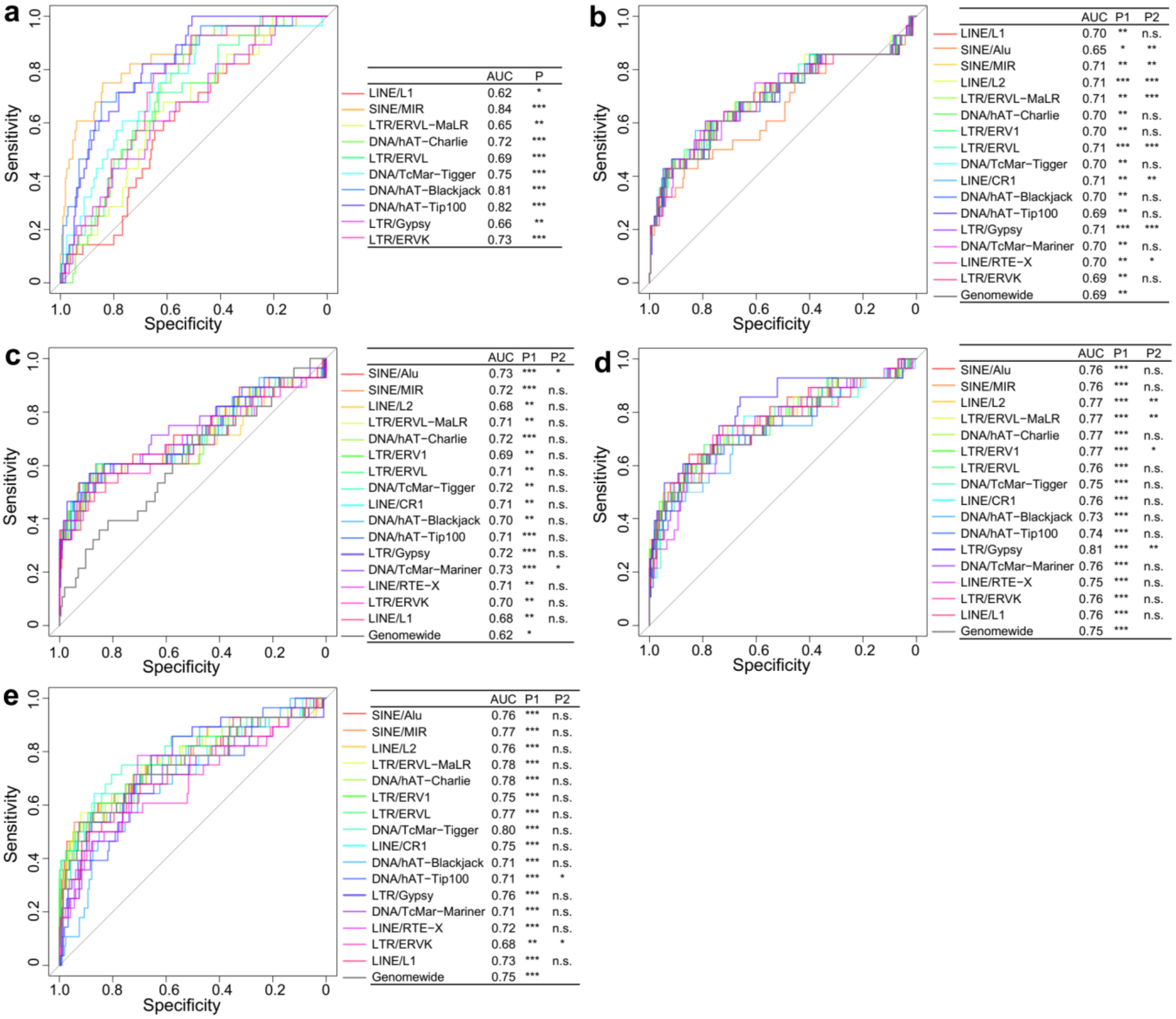


**Fig. S17. ROC curves for TEs to differentiate ovarian cancer samples from controls in Cristiano et al. dataset.** Receiver Operating Characteristic (ROC) curves for (a) RSD values, (b) E-index values, (c) short cfDNA fragments, (d) end motif diversity score, and (e) CCCA end motif usages in TEs and genomewide level to differentiate bile duct cancer samples from controls. P1 represented p-value for the AUC calculated using Z-tests, and P2 represented p-value comparing the ROCs of TE versus genomewide level using DeLong tests. *p<0.05, **p<0.01, ***p<0.001.

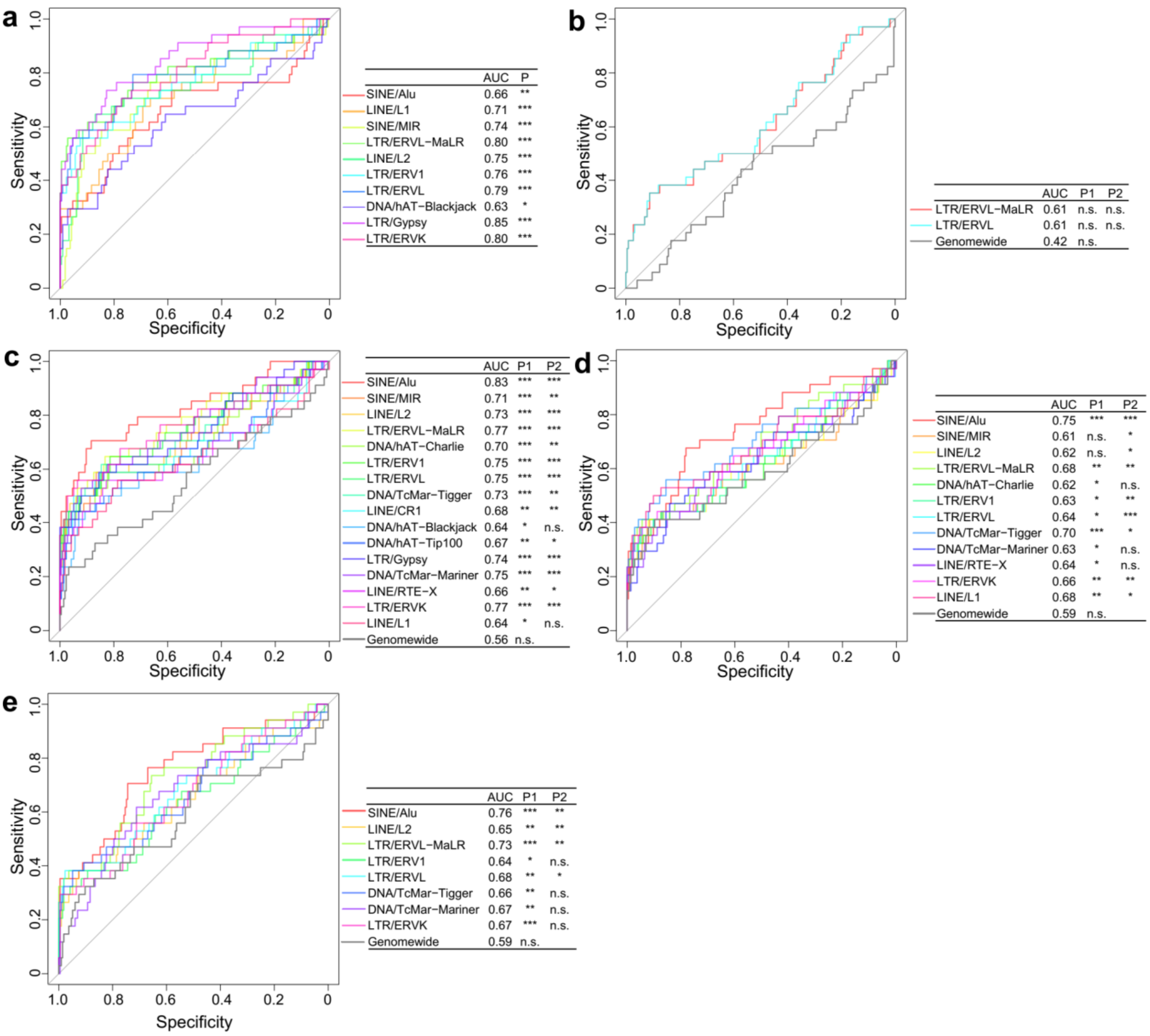


**Fig. S18. ROC curves for TEs to differentiate pancreatic cancer samples from controls in Cristiano et al. dataset.** Receiver Operating Characteristic (ROC) curves for (a) RSD values, (b) E-index values, (c) short cfDNA fragments, (d) end motif diversity score, and (e) CCCA end motif usages in TEs and genomewide level to differentiate bile duct cancer samples from controls. P1 represented p-value for the AUC calculated using Z-tests, and P2 represented p-value comparing the ROCs of TE versus genomewide level using DeLong tests. *p<0.05, **p<0.01, ***p<0.001.

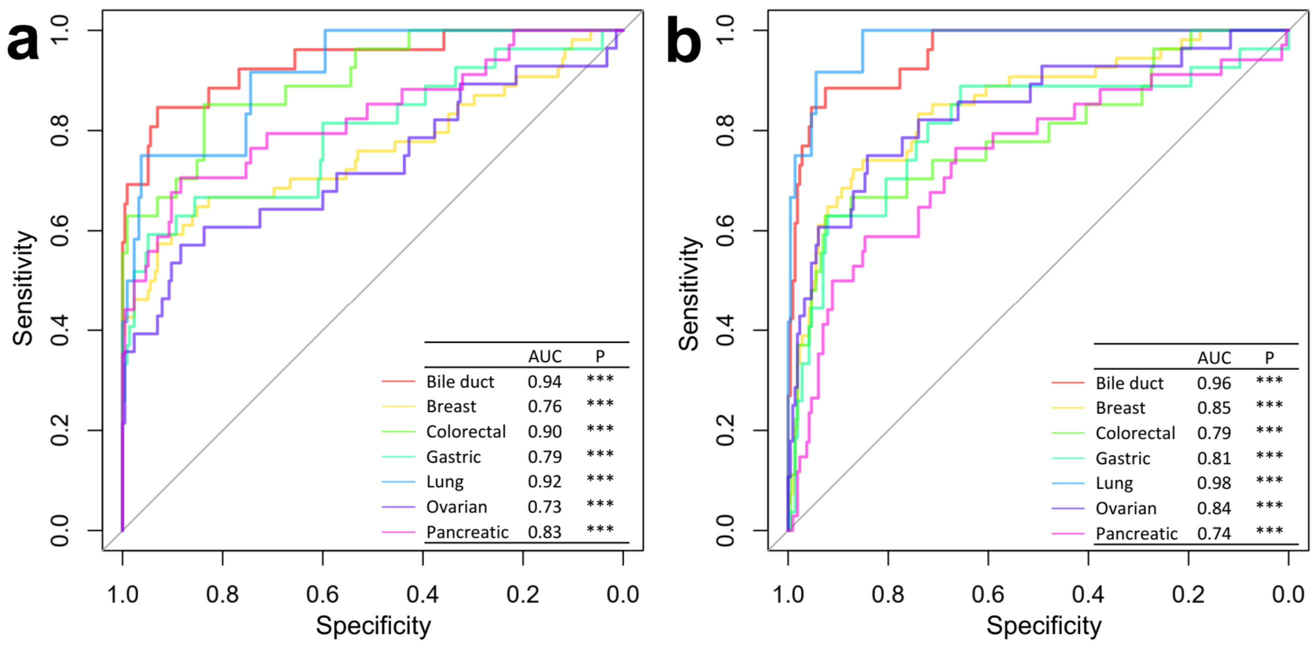


**Fig. S19. ROC curves for (a) using E-index values in SINE/ALU to differentiate cancer samples from controls, and (b) using RSD values in SINE/MIR to differentiate cancer samples from controls in Cristiano et al. dataset.** P-values for AUCs were calculated using Z-tests. ***p<0.001.

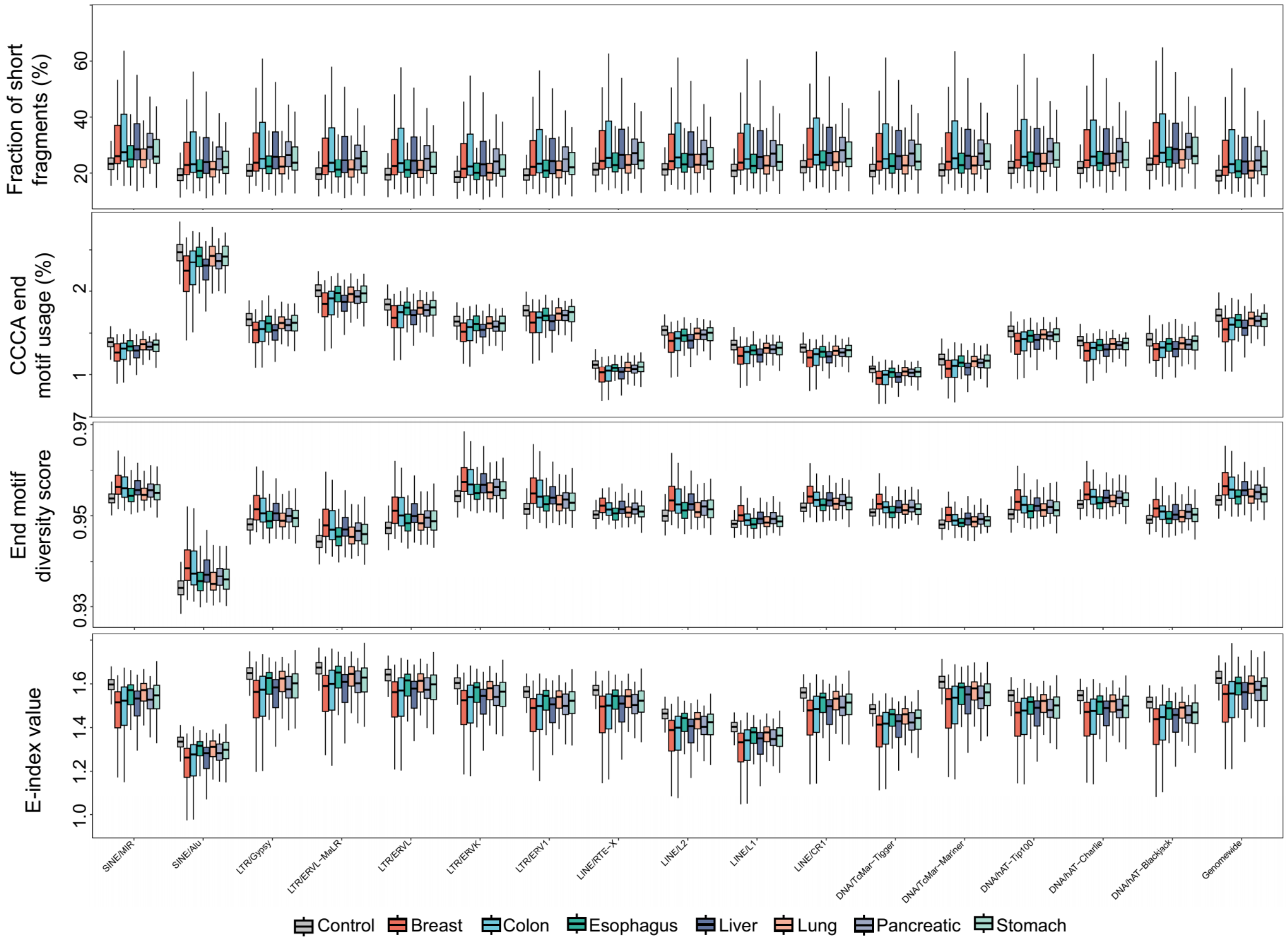


**Fig. S20. CfDNA fragmentomic characteristics in TEs in Bie et al. dataset.** Fraction of short cfDNA fragments (i.e., ≤ 150bp), motif patterns, and E-index values between controls and cancer patients across various TEs. Grey, red, blue, green, dark purple, orange, light purple, and light green boxes presented controls, breast cancer, colorectal cancer, esophagus cancer, liver cancer, lung cancer, pancreatic cancer, and stomach cancer samples, respectively.

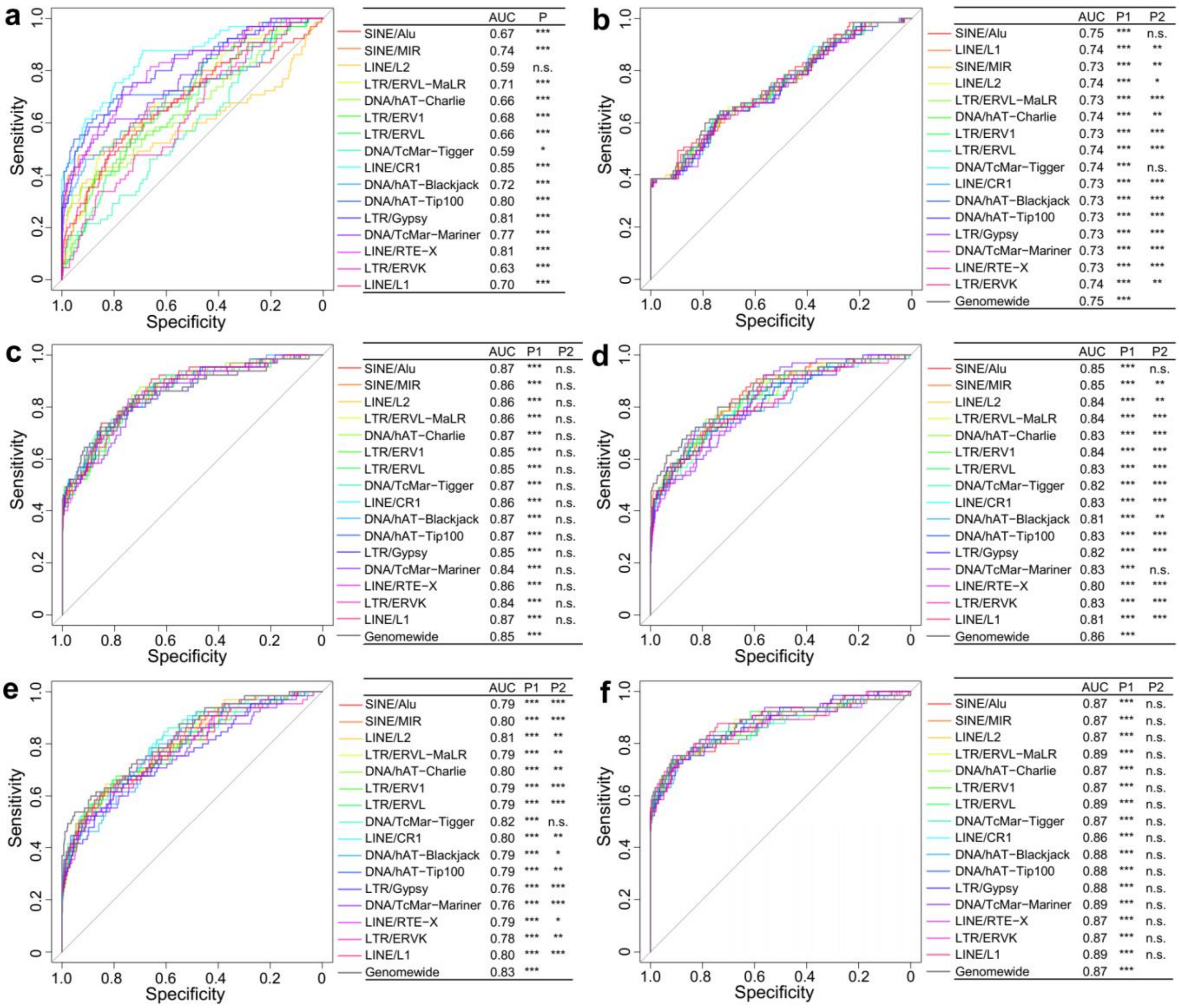


**Fig. S21. ROC curves for TEs to differentiate breast cancer samples from controls in Bie et al. using (a) RSD values (b) fraction of short cfDNA fragments, (c) E-index values, (d) end motif diversity score, (e) CCCA end motif usages, and (f) DNA methylation levels.** P and P1 represented p-values for AUCs calculated using Z-tests, and P2 represented p-values comparing the ROCs of each TE versus genomewide level using DeLong tests. *p<0.05, **p<0.01, ***p<0.001.

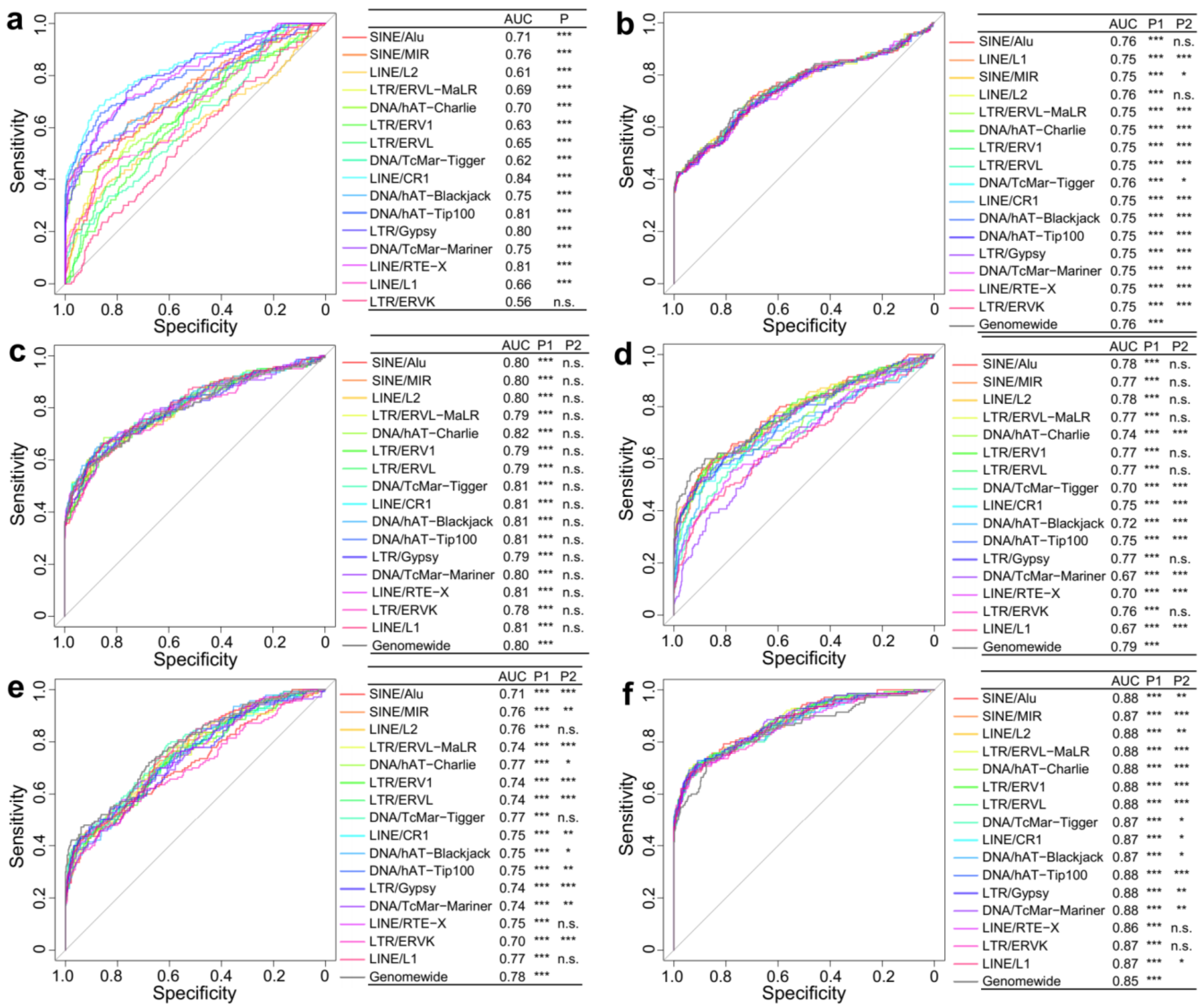


**Fig. S22. ROC curves for TEs to differentiate colorectal cancer samples from controls in Bie et al. using (a) RSD values (b) fraction of short cfDNA fragments, (c) E-index values, (d) end motif diversity score, (e) CCCA end motif usages, and (f) DNA methylation levels.** P and P1 represented p-values for AUCs calculated using Z-tests, and P2 represented p-values comparing the ROCs of each TE versus genomewide level using DeLong tests. $*p<0.05$, $**p<0.01$, $***p<0.001$.

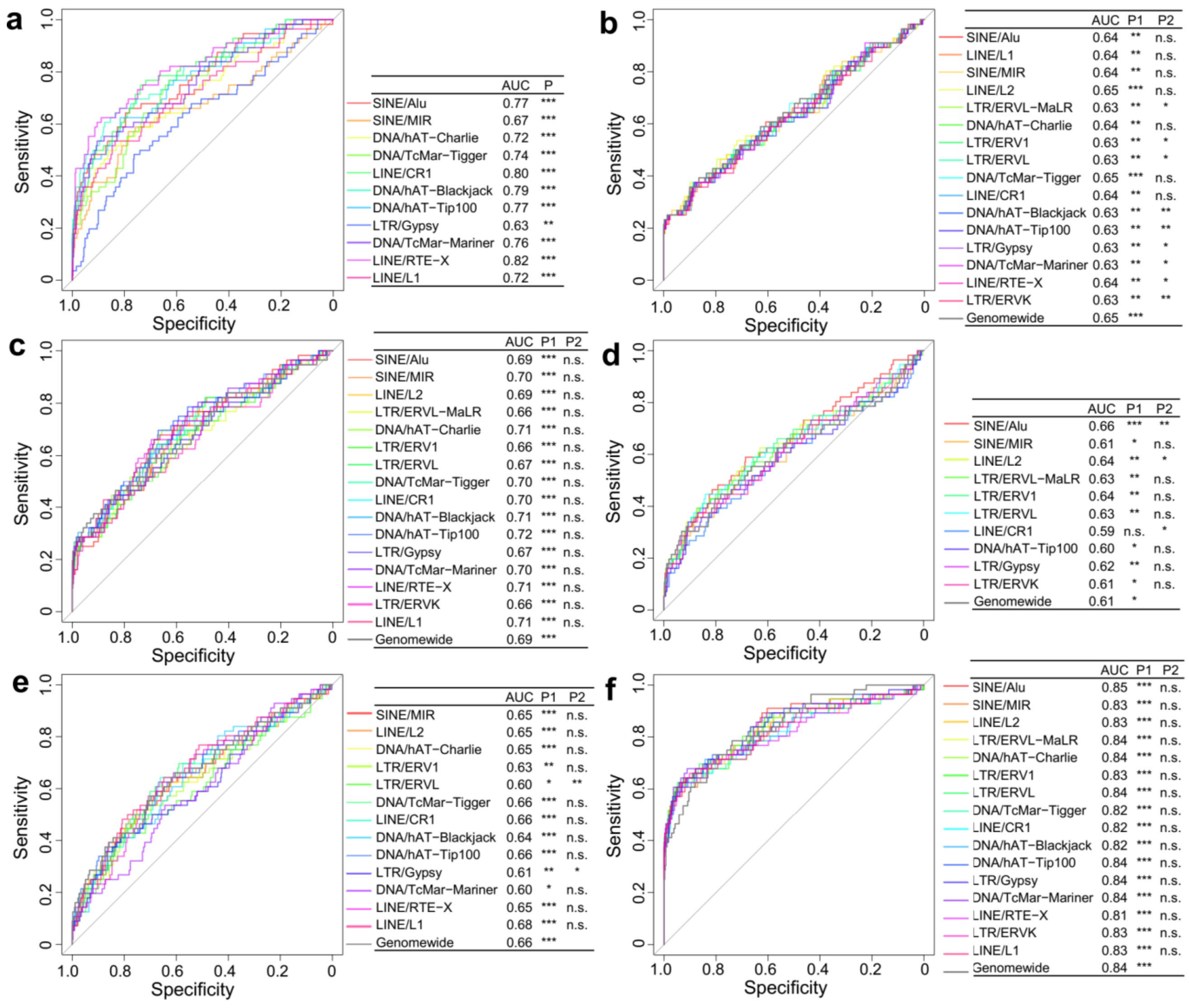


**Fig. S23. ROC curves for TEs to differentiate esophagus cancer samples from controls in Bie et al. using (a) RSD values (b) fraction of short cfDNA fragments, (c) E-index values, (d) end motif diversity score, (e) CCCA end motif usages, and (f) DNA methylation levels.** P and P1 represented p-values for AUCs calculated using Z-tests, and P2 represented p-values comparing the ROCs of each TE versus genomewide level using DeLong tests. *p<0.05, **p<0.01, ***p<0.001.

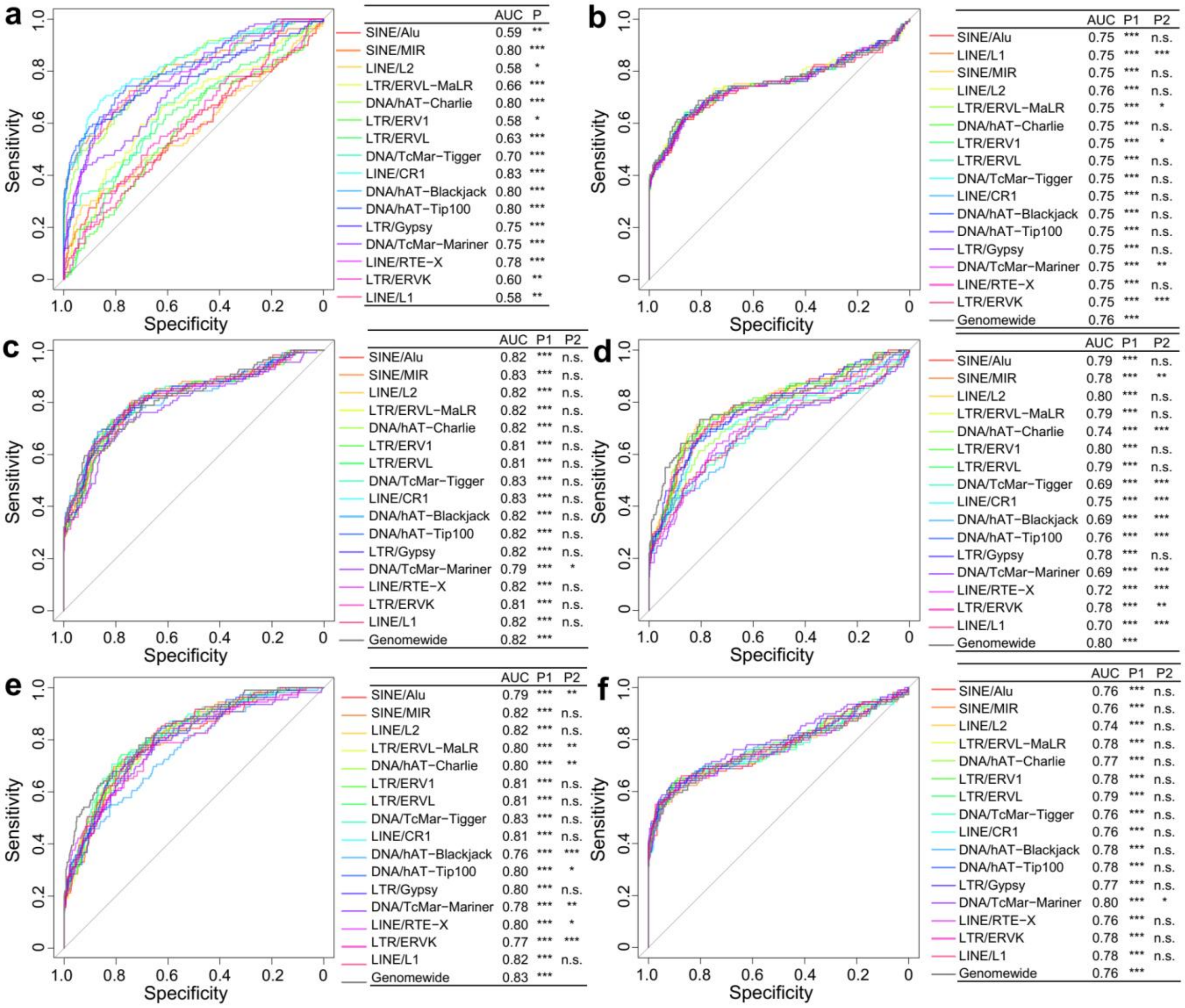


**Fig. S24. ROC curves for TEs to differentiate liver cancer samples from controls in Bie et al. using (a) RSD values (b) fraction of short cfDNA fragments, (c) E-index values, (d) end motif diversity score, (e) CCCA end motif usages, and (f) DNA methylation levels.** P and P1 represented p-values for AUCs calculated using Z-tests, and P2 represented p-values comparing the ROCs of each TE versus genomewide level using DeLong tests. *p<0.05, **p<0.01, ***p<0.001.

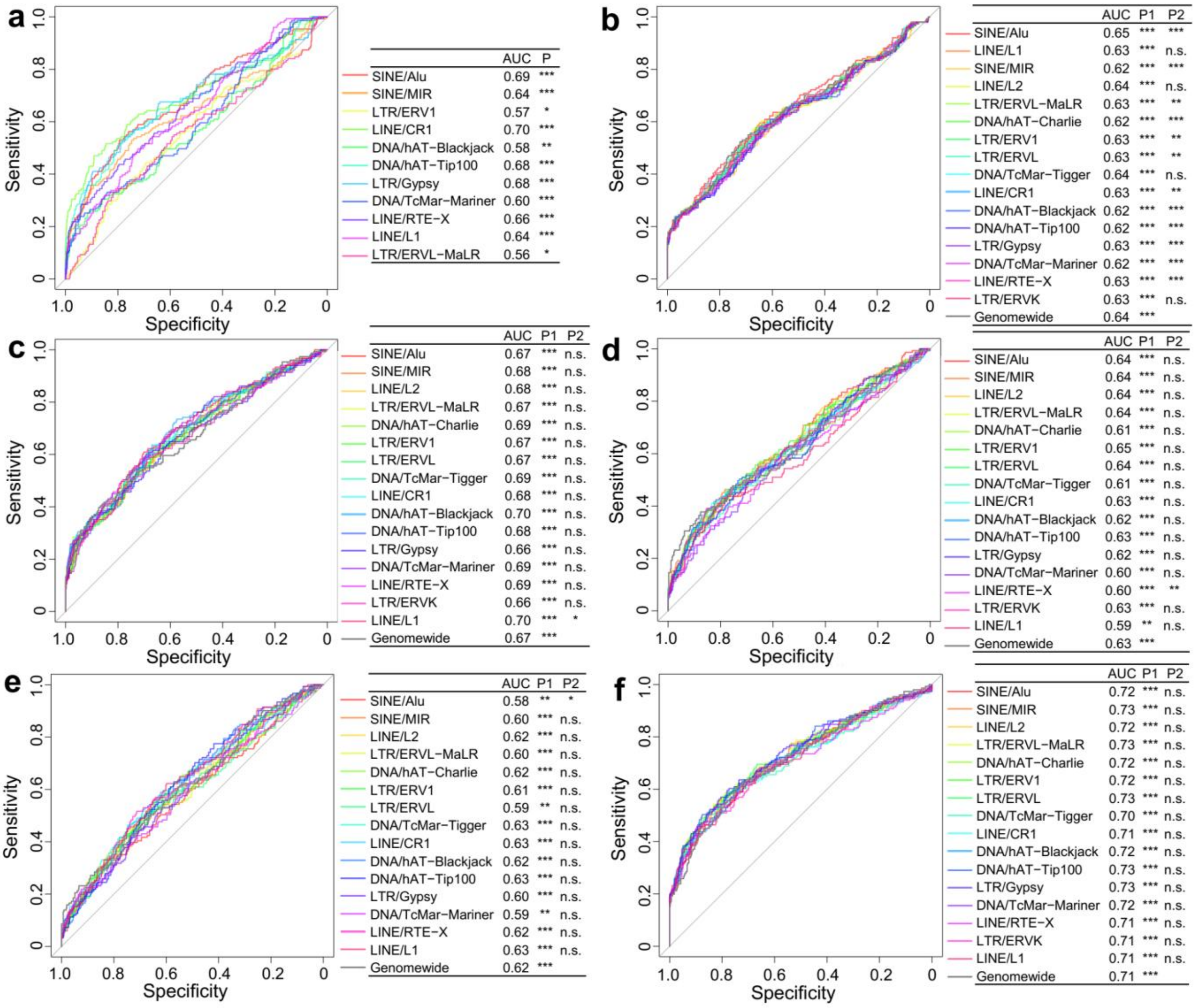


**Fig. S25. ROC curves for TEs to differentiate lung cancer samples from controls in Bie et al. using (a) RSD values (b) fraction of short cfDNA fragments, (c) E-index values, (d) end motif diversity score, (e) CCCA end motif usages, and (f) DNA methylation levels.** P and P1 represented p-values for AUCs calculated using Z-tests, and P2 represented p-values comparing the ROCs of each TE versus genomewide level using DeLong tests. *p<0.05, **p<0.01, ***p<0.001.

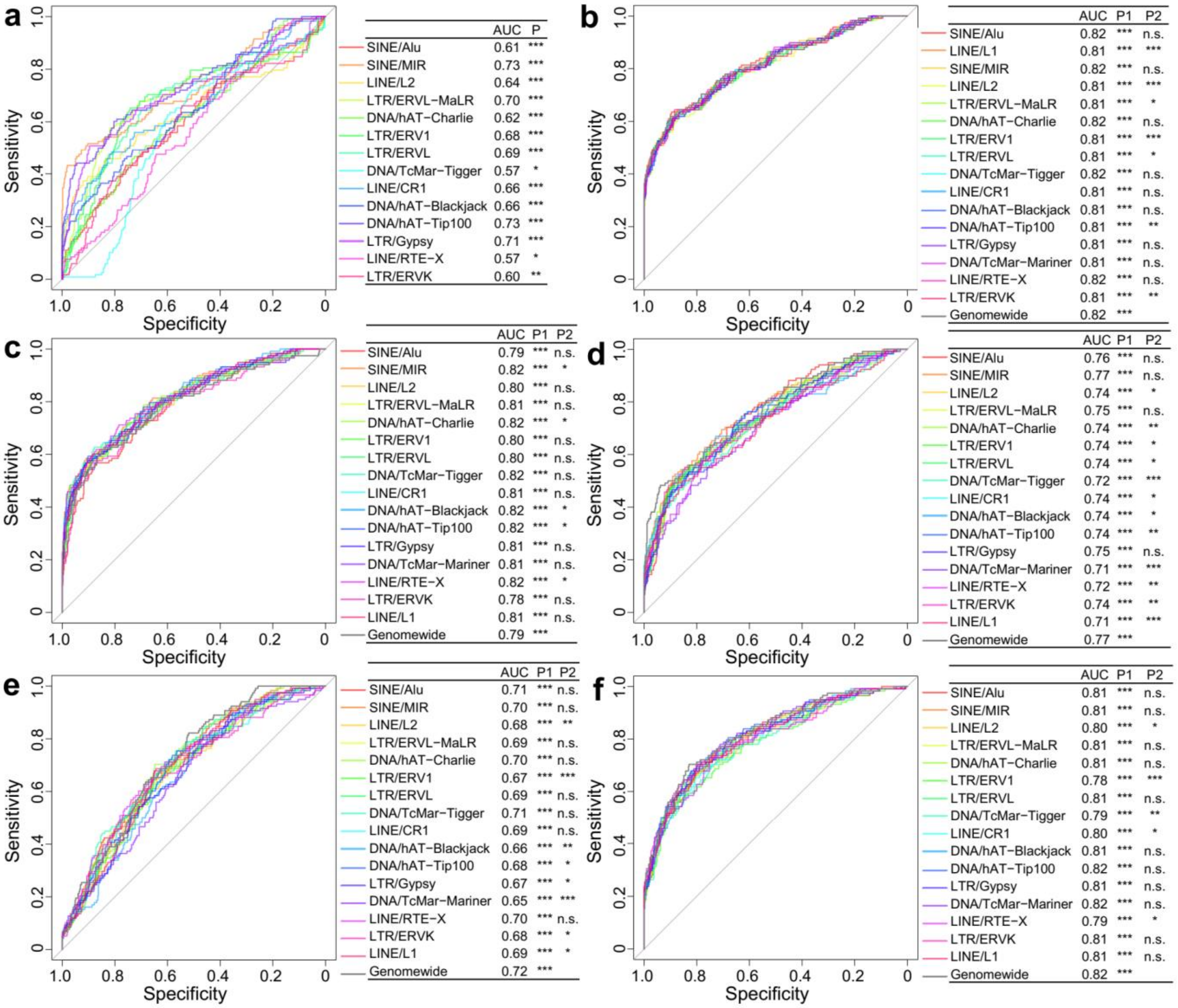


**Fig. S26. ROC curves for TEs to differentiate pancreatic cancer samples from controls in Bie et al. using (a) RSD values (b) fraction of short cfDNA fragments, (c) E-index values, (d) end motif diversity score, (e) CCCA end motif usages, and (f) DNA methylation levels.** P and P1 represented p-values for AUCs calculated using Z-tests, and P2 represented p-values comparing the ROCs of each TE versus genomewide level using DeLong tests. *p<0.05, **p<0.01, ***p<0.001.

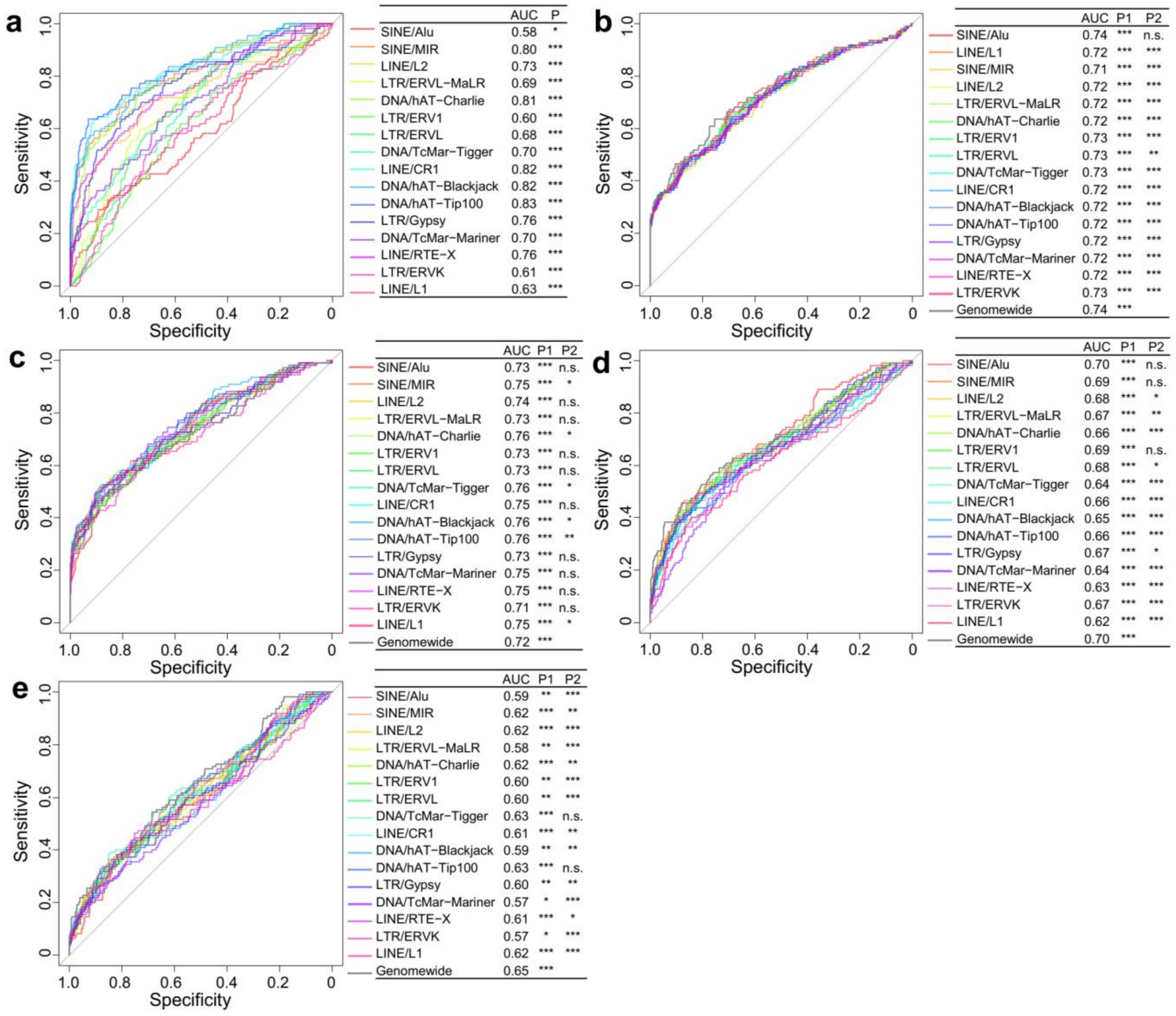

**Fig. S27. ROC curves for TEs to differentiate stomach cancer samples from controls in Bie et al. using (a) RSD values (b) fraction of short cfDNA fragments, (c) E-index values, (d) end motif diversity score, and (e) CCCA end motif usages.** P and P1 represented p-values for AUCs calculated using Z-tests, and P2 represented p-values comparing the ROCs of each TE versus genomewide level using DeLong tests. *p<0.05, **p<0.01, ***p<0.001.
